# Signatures of Micropeptides Encoded by lncRNAs in Cancer Progression and Metastasis

**DOI:** 10.1101/2025.09.22.677872

**Authors:** Stav Zok, Michal Linial

**Affiliations:** The Rachel and Selim Benin School of Computer Science and Engineering, The Hebrew University of Jerusalem, Jerusalem, Israel; Department of Biological Chemistry, The Life Science Institute, The Hebrew University of Jerusalem, Israel

**Keywords:** TCGA, AlphaFold, epigenetic regulation, signal peptide, antisense, Ribo-seq, LncBook, SNHG

## Abstract

Long non-coding RNAs (lncRNAs) are key regulators of gene expression, chromatin remodeling, and signaling. Recent estimates suggest that the human genome contains more than 35,000 lncRNA genes, with roughly 20% predicted to encode micropeptides (MPs) with unknown functions. In this study, we focused on the subset of lncRNAs with strong statistical evidence for MP-encoding potential, accounting for approximately 8% of the unfiltered MPs collection. Our analysis centered on 1,782 high-confidence lncRNA-MPs derived from 478 genes expressed across 17 cancer types from The Cancer Genome Atlas (TCGA). We show that lncRNA-MPs display distinct amino acid compositions and unique 4-mer patterns compared to the human coding proteome. A few genes (9) with exceptionally long transcripts are characterized by ≥20 MPs each. Functional interference confirmed that most of the lncRNA-MPs are unstructured. Only a third of the genes display some phylogenetic conservation, and only 4 genes display canonical N-terminal signal peptides characteristic of secreted proteins. We focused on cancer progression-associated lncRNAs that show differential expression (z-score >|3|) across consecutive tumor stages and metastatic states (transitional lncRNAs, Tr-lncRNAs). A collection of 72 genes encoding 314 MPs (Tr-lncRNA-MPs) was detected, with 76% of the MPs being ≥30 amino acids long. Prediction by AlphaFold 2.0 and homology modeling tools revealed dozens of MPs with well-defined secondary structures and recognizable 3D motifs. Among the longer Tr-lncRNA-MPs (>60 amino acids), we confirmed the presence of ubiquitin-like, RNase H-related, and other conserved foldable motifs. Known cancer lncRNAs containing high-confidence MPs (XIST, UCA1, HOXA11-AS, LINC01234, and HAND-AS1) overlap with 50 pan-cancer lncRNAs associated with tumor stage or metastasis transitions. Together, these findings demonstrate that integrating sequence motifs (e.g., signal peptides, k-mers) with structural foldability offers a multifaceted view of lncRNA-MPs in cancer. We argue that the capacity to produce MPs may reinforce the oncogenic impact dominated by the lncRNA entity. We propose that Tr-lncRNA-MPs represent a promising new class of biomarkers and therapeutic targets in oncology.

**Key points:**

- 478 lncRNA genes with strong evidence for micropeptide (MPs) production generated 1,782 distinct lncRNA-MPs.
- 72 lncRNAs and 314 MPs are associated with transitional lncRNAs from 17 cancer types and stages of tumor progression and metastasis.
- Sequence and structural analyses reveal many MPs with reliable 3D folding potential.
- Dozens of previously overlooked MPs may serve as novel biomarkers and therapeutic targets in cancer.

## Introduction

Cataloging long non-coding RNA (lncRNAs) identified over 35k genes and approximately 190k transcripts in the annotated human genome ^1^. While the regulation of lncRNA expression is similar to that of coding mRNAs, lncRNAs possess distinct features. Most lncRNAs contain a small number of exons (2 on average), are poorly conserved, and predominantly localize to the nucleus and chromatin. Based on expression analysis, most lncRNAs exhibit low expression levels but are characterized by high tissue and cell specificity ^2^. Advances in proteomics and translation technology have led to the discovery of MPs encoded by short ORFs (sORFs) within lncRNA sequences, typically ranging from 10 aa to 100 aa ^3^. Multiple ribosome-profiling and proteogenomic studies suggest that hundreds to a few thousand putative micropeptides (MPs) may be produced from lncRNAs. However, high-confidence, experimentally supported cases are sparse, and strong evidence is limited to only tens of genes ^4^. Some of these lncRNA-MPs act as tumor suppressors, while others promote cancer growth by influencing key biological processes. The question of what constitutes sufficient evidence for an lncRNA to be truly “coding” remains unresolved ^4^.

Ribosome binding alone, according to Ribo-seq, does not necessarily confirm the production of a functional peptide ^5^. Therefore, studying lncRNA-encoded peptides is challenging, and support for their coding capacity relies on a combination of computational and experimental approaches ^6, 7^. The relatively poor evolutionary conservation of lncRNAs reduces the utility of cross-species conservation tools (e.g., PhyloCSF) for predicting whether an sORF is likely to code for a functional peptide ^8^. Like canonical proteins, lncRNA-MPs generally begin with a methionine (Met) and are transcribed as capped and polyadenylated transcripts that are exported to the cytoplasm ^9^. However, it has been shown that under stress conditions, translation from lncRNAs may be initiated without a cap or classical ribosome binding sites. Instead, an internal ribosome entry site (IRES) may be used to initiate translation ^10^.

A question about how many proteins might be produced from non-canonical ORFs derived from lncRNAs remains open (see estimated in ^5^). Direct detection methods like antibody-based assays and MS provide evidence by identifying the actual peptides. Surprisingly, Ribo-seq data in mouse cells have shown that almost half of lncRNAs may contain translated small ORFs (sORFs). However, direct mass spectrometry (MS) peptidomics, combined with RNA-seq from human K562 cells, identified only minute amounts of small peptides. Moreover, from the MS results, it was confirmed that many identified sORFs lack Met as a start codon ^11^. PepScore provides a comprehensive analysis of genome-wide translated ORFs using ribosome profiling data ^12^. The score aims to predict the likelihood that a given peptide is stable in human cells. This study showed that inhibiting proteasomal or lysosomal degradation could stabilize some moderately scored peptides, while low-scoring ones are often rapidly degraded by proteases ^12^.

Until recently, most findings and experimental evidence on open reading frames (ORFs) from long non-coding RNAs (lncRNAs) were limited to specific examples. A few dozen lncRNA-encoded MPs, typically 30-100 amino acids (aa) in length, have been experimentally validated in the context of specific cancer types ^13^. However, their specific biological functions remain largely unknown. One example is the HOXB-AS3 MP, a short peptide (53 aa) that acts as a tumor suppressor in colon cancer by attenuating aerobic glycolysis. Specifically, the peptide blocks the hnRNPA1-mediated splicing of pyruvate kinase M (PKM) mRNA. Interestingly, the selection of the alternatively spliced PKM variant is associated with the Warburg effect and tumor growth. The loss of HOXB-AS3 MP correlates with poor prognosis and enhanced metabolic reprogramming ^14^. In contrast, HOXB-AS3 overexpression in OSCC is associated with poor patient prognosis, where it directly interacts with the RNA-binding protein IGF2BP2, leading to stabilization of the c-Myc oncogene that drives cancer cell growth ^15^.

The established role of lncRNA LINC00992 was attributed to its MP called GT3-INCP (GATA3-interacting cryptic protein; 120 aa). GT3-INCP promotes estrogen receptor-positive (ER+) breast cancer progression ^16^. In this case, the lncRNA-derived peptide directly binds to GATA3, a lineage-defining transcription factor (TF), and facilitates hormone-driven cancer transition. The presence of GT3-INCP enables cancer cells to escape hormone dependency therapy in breast cancer patients. A recent review article provides a detailed summary of a couple of dozen MPs and their effects in specific cancer types and cellular contexts ^6^. MPs might be used in medical applications, potentially as diagnostic markers or therapeutic targets ^17, 18^.

In this study, we aim to characterize the protein capacity of the lncRNAs and their occurrence in major cancer types, using Ribo-seq profiling data from SmProt and TCGA data for the cancer angle. The specificity of lncRNAs was especially evident in cancer, as revealed by analyzing the lncRNA expression from TCGA ^19^. Many lncRNAs that are differentially expressed across transitions are unique to specific cancer types. We focused on the subset of lncRNAs whose expression shows significant changes along transitions (named Tr-lncRNAs). To this end, we characterized the lncRNAs with confident protein capacity, based on MPs that were assigned with statistically significant experimental evidence. The nature of the lncRNAs-MPs was reported for the 1,782 high-confidence peptides, according to AlphaFold2.0 structural prediction ^20^. A pan-cancer view yielded insights into the set of features that specify lncRNA encoded MPs across cancer types. We discuss the potential role of lncRNA-MPs in diagnosis, prognosis, and as a lead to activating the immune system and in cell-cell communications along cancer progression and metastasis.

## Results

### Most lncRNAs are cancer transition-specific

Long non-coding RNAs (lncRNAs) are broadly defined as non-coding transcripts longer than 200 nucleotides. In cells, lncRNAs act via multiple, non-exclusive modes of action, including peptide production. From a collection of 190k transcripts associated with lncRNAs (GENECODE V49), we focus on a small subset of lncRNAs that are likely to act via their micropeptides (MPs) in cancer samples from major cancer types from TCGA. We compiled a set of 1,782 (Supplementary **Table S1**) that was derived from 478 lncRNA genes, which are considered high-confidence MPs (lncRNA-MPs). Note that this collection represents 8% of the unfiltered collection of potential peptides (see Methods).

We focused on lncRNAs that may be acting in cancer transition. To this end, we compiled the public data from the TCGA with lncRNA-mapped genes and performed two main sets of analyses: Tumor stage-based (T-stage) and Metastatic-based (M-state). Genome coordinates were used to confirm unique mapping, resulting in a final gene list of 13,257 lncRNA genes. We analyze each of these lncRNAs according to their differential expression behavior for any consecutive stage (partitioned to up- and downregulation), and across major cancer types. We observed many lncRNAs that are associated with a specific cancer type and a unique transition. The lncRNAs that are differentially expressed (z-score >|3|) and exhibit consistency across cancer samples are termed transitional lncRNAs (Tr-lncRNAs, ^21^). The tumor transitions by stages (T-stage) include I→II, II→III, III→IV, and a simplified combination of I and II and III and IV, marked as E→L, for early to late tumor progression. In terms of metastasis state (M-state), we determine the transition as a non-metastasis sample marked by the tumor size, categorized as small and large (M0s→M0l), transition to metastasis indicated as M0l→M1, and simplified category of ME→ML (early to late). Altogether, there are 7 transitions (note that E→L transitions are used for coarse partition of cancer samples).

Supplementary **Fig. S1** illustrates Upset plots for the upregulated and downregulated lncRNAs for the E❼L transition (marked positive and negative, respectively). Inspection of the plots shows that the vast majority of Tr-lncRNAs are specific to only one cancer type and mostly for one transition (analyzed in ^21^). For example, there are 40 of 44 Tr-lncRNAs that are uniquely associated with the downregulation of E❼L in UCEC (Supplementary **Fig. S1**). Our analysis covers 17 cancer types represented in TCGA (>150 samples each). Altogether, for all 7 transitions tested (4 for T-stage and 3 for M-state), we identified 2,399 Tr-lncRNAs ^21^. Supplementary **Table S2** lists the TCGA abbreviations of cancer types along with basic statistics.

### A small fraction of Tr-lncRNAs encode micropeptides (MP)

In this study, we focus on the protein capacity from lncRNAs (lncRNA-MPs), and specifically on the subset that is represented in cancer samples, and among them, the set of transition-specific (Tr-lncRNAs). **Fig. 1** shows Tr-lncRNAs in major cancer types for T-stage transition E❼L. The bars for each of the 16 major cancer types show the fraction of unfiltered lncRNA-MPs among Tr-lncRNAs along with their statistical enrichment (**Fig. 1A**). For example, the average fraction of lncRNA-MPs across all cancer types is ∼21%, but it accoun for 55% in the COAD positive expression trend of E❼L transition. The fractions of the listed MPs in eac cancer type are statistically significant (corrected p-value <0.05, excluding KIRC).

**Fig. 1.**
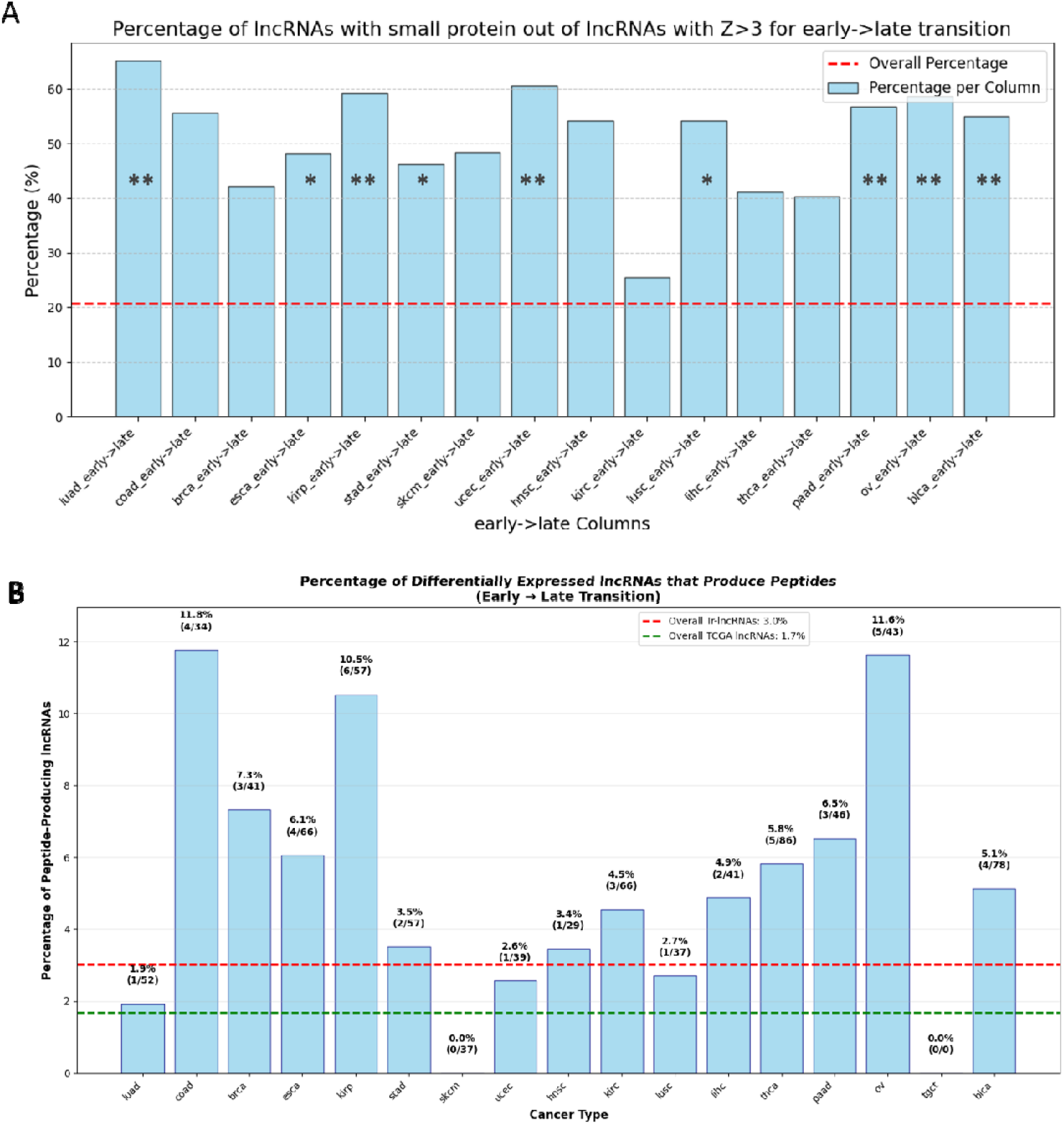
The percentage of MPs containing lncRNAs among Tr-lncRNAs across cancer types. **(A)** The fraction of E❼L transition among 16 major cancer types. Only upregulated sets for E❼L transition are shown for any annotated MPs. The dashed red horizontal line reflects the average percentage of MPs containing lncRNAs among all lncRNAs. Corrected p-value <1e-6 and <1e-4 are marked by ** and *, respectively. The fraction of MPs in each cancer type (excluding KIRC) is significant with a corrected p-value <0.05. **(B)** The fraction of E❼L transition among major cancer types for upregulated sets. Only Tr-lncRNAs with confident MPs are considered. The number of cases is indicated above the bars. Statistical analysis was not performed due to the small number of instances (total 72 confident Tr-lncRNA-MPs). The dashed green and red horizontal lines represent the average percentage of confident MPs among all lncRNAs and among the Tr-lncRNAs, respectively.

We further filtered out Tr-lncRNA by demanding their peptide capacity. We applied a confidence threshold using the internal statistics of the SmProt scoring. The SmProt considers high confidence MP by matching with TIS and the properties of the experimental results from Ribo-seq (see Methods). **Fig. 1B** represents the reduced number of Tr-lncRNAs for the strongly supported lncRNA-MPs. Note that by selecting Tr-lncRNA-MPs with strong evidence, the fraction of lncRNA-MPs that was 21% in the unfiltered set (**Fig. 1A**) was only 3% among the high-quality MPs (**Fig. 1B**). Still, we show that the overall frequency of high confidence Tr-lncRNA-MPs is significantly higher than the default occurrence of lncRNA-MPs (**Fig. 1B**, dashed green line, accounting for 1.7% out of all TCGA lncRNA genes). Strongest enrichment of lncRNA-MPs (i.e., >10% of the reported Tr-lncRNA) is linked to COAD, KIRC, and OV cancers. Restricting candidate Tr-lncRNA genes to a small, high-confidence set of MPs results in analyzing most reliable lncRNAs, while simultaneously requiring evidence of their functional relevance to cancer progression.

Altogether for any of the transitions, cancer types, and expression directionality (up- and downregulation), we report on 478 lncRNAs that are the source of 1,782 peptides (≥10 amino acids; Supplementary **Table S1**). The rest of the analysis will refer to this strict set of genes and their respective peptides.

### The protein capacity of lncRNAs in TCGA

Despite the definition of lncRNAs as non-coding entities, emerging studies reveal the importance of lncRNA-encoded micropeptides (MP) in oncogenesis ^13^. A handful of validated examples of mesenchymal transition (EMT), metastasis, and treatment resistance. **Fig. 2A** presents the length distribution of lncRNA-MPs. It shows that the length of the MPs is inversely proportional to the amounts, as expected for statistical candidates of ORFs from lncRNAs. The distribution of MP length for lncRNAs that are involved in cancer transition (according to differentially expressed |z-values|>3) resulted in a substantial shift toward longer MPs (60% relative to 52% of MP>30 aa; **Fig. 2B)**. Repeating the peptide length analysis for the strict collection of 1,782 peptides (478 lncRNA-MPs) shows that while the amount of confident lncRNA-MPs is less than 10% of the the unfiltered lncRNA-MPs, the length profile is maintained for all lncRNA-MPs (**Fig. 2C**) and for the Tr-lncRNA-MPs (**Fig. 2D**). The following analyses are restricted to the confident lncRNA-MPs. **Fig. 1E** shows the distribution of the confident MP counts per lncRNA gene. The number of MPs per lncRNA for lncRNAs that have ≥4 MP is substantial 28.8% and 32%, considering the lncRNA-MP and the Tr-ncRNA-MP sets, respectively (**Fig. 1D**).

**Fig. 2.**
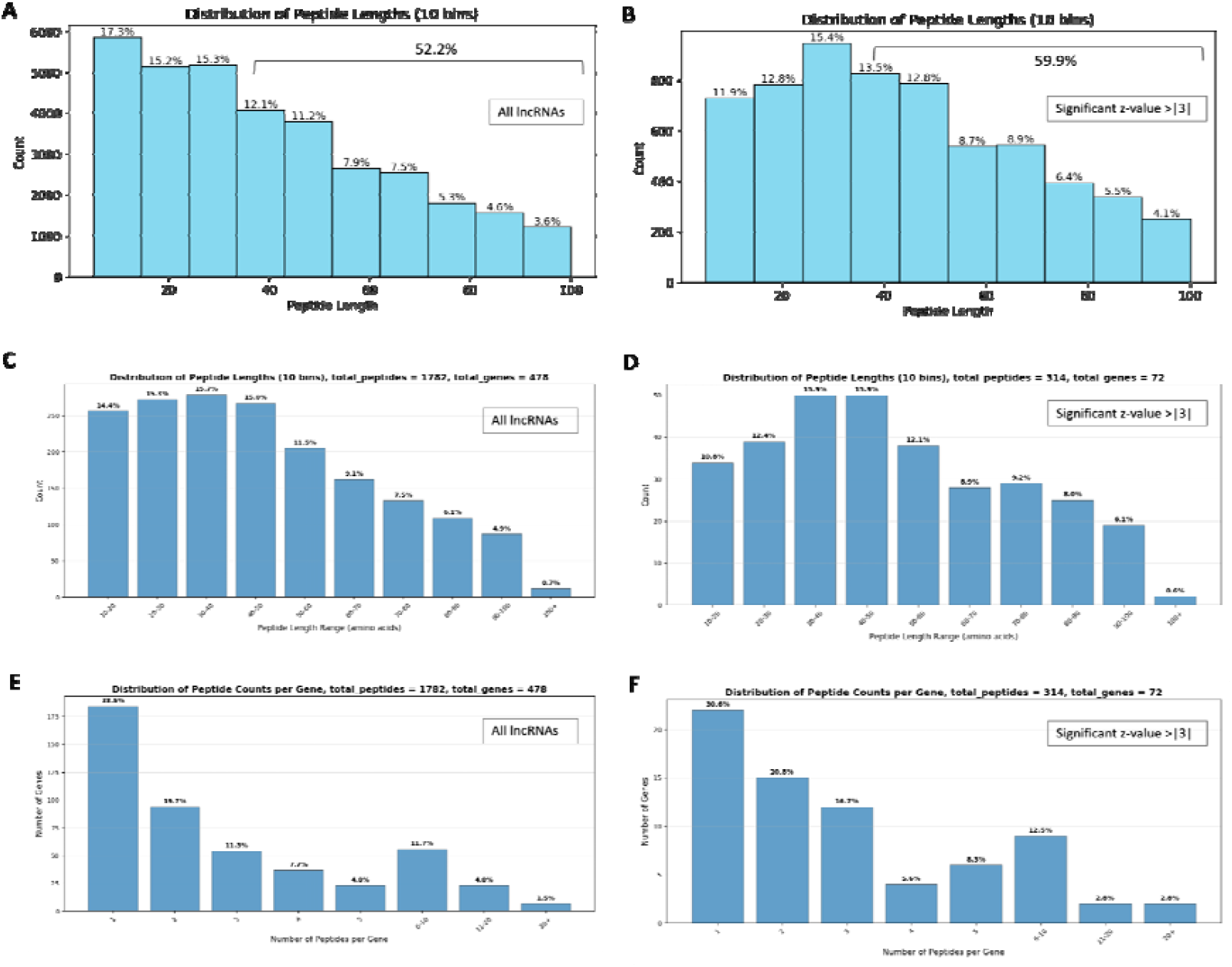
Properties of lncRNA-MPs**. (A)** Distribution of peptide length of all lncRNAs by 10 bins. The lengths of 52.2% of all lncRNA-MPs are ≥30 aa. The longest peptides were 100 aa in length. **(B)** Distribution of peptide length of all lncRNAs that have a significant |z-value| > 3. The lengths of 59.9% of all lncRNA-MPs are ≥30 aa. **(c)** Distribution of peptide length of all confident lncRNA-MPs. The number of peptides and genes is indicated. The longest peptides were 100 aa in length. **(D)** Distribution of peptide length of all lncRNAs that have a significant |z-value|> 3 (i.e., Tr-lncRNA-MPs). The number of peptides and genes is indicated. **(E)** The percentage of MPs partitioned by bins for the number of MPs for each lncRNA for all lncRNAs.**(F)** The percentage of MPs is partitioned into bins based on the number of MPs per Tr-lncRNA group. The number of peptides and genes is indicated.

### Only a third of the confident MP candidates start with initiating methionine

Translation of the human proteome is expected to initiate with methionine (Met, M) as part of the Kozak ribosomal binding site (ATG codon, **Fig. 3A**). In the context of sORFs and lncRNA-MPs, initiation may occur at near-cognate start sites. The data that was based on Ribo-seq provides some evidence for the non-ATG initiation of translation. A single base change of ATG is still able to initiate translation (i.e., TTG, GTG, CTG, AAG, AGG, ACG, ATA, ATT, and ATC; **Fig. 3A**). The occurrence of near-cognate sites is in accord with initiation by Leu, Val, Lys, Arg, Thr and Ile (**Fig. 3A**). **Fig. 3B** shows the confident set of MPs (>10 aa), indicating that only 30% of the peptides start with Met. This ratio was quite insensitive to the peptide length or to the subset of Tr-lncRNA-MPs. Among the Tr-lncRNA-MPs, approximately half of the peptides initiated with either Met or Leu (**Fig. 3B**, right). This is a significantly lower fraction relative to the results from analyzing the unfiltered lncRNA-MPs. Upon removal of redundancy (by selecting a representative for a cluster sharing >90% aa sequence identity), this fraction increased to 37.7% (**Fig. 3C**, left). Interestingly, this value remained stable for the Tr-lncRNA-MPs, which were clustered by Ref90 and the more stringent threshold of Ref60 protocols (**Fig. 3C**). We conclude that MPs originating from Tr-lncRNAs tend to use Met only slightly more with respect to the collection of all high-confidence lncRNA-MPs. We also observed that two-thirds of the MP representatives (**Fig. 3C**, right) initiated with either Met or Leu. We concluded that the near-cognate initiation should be considered with preference to Met and Leu, and along with a far lower preference to K, R, and T as initiating codons.

**Fig. 3.**
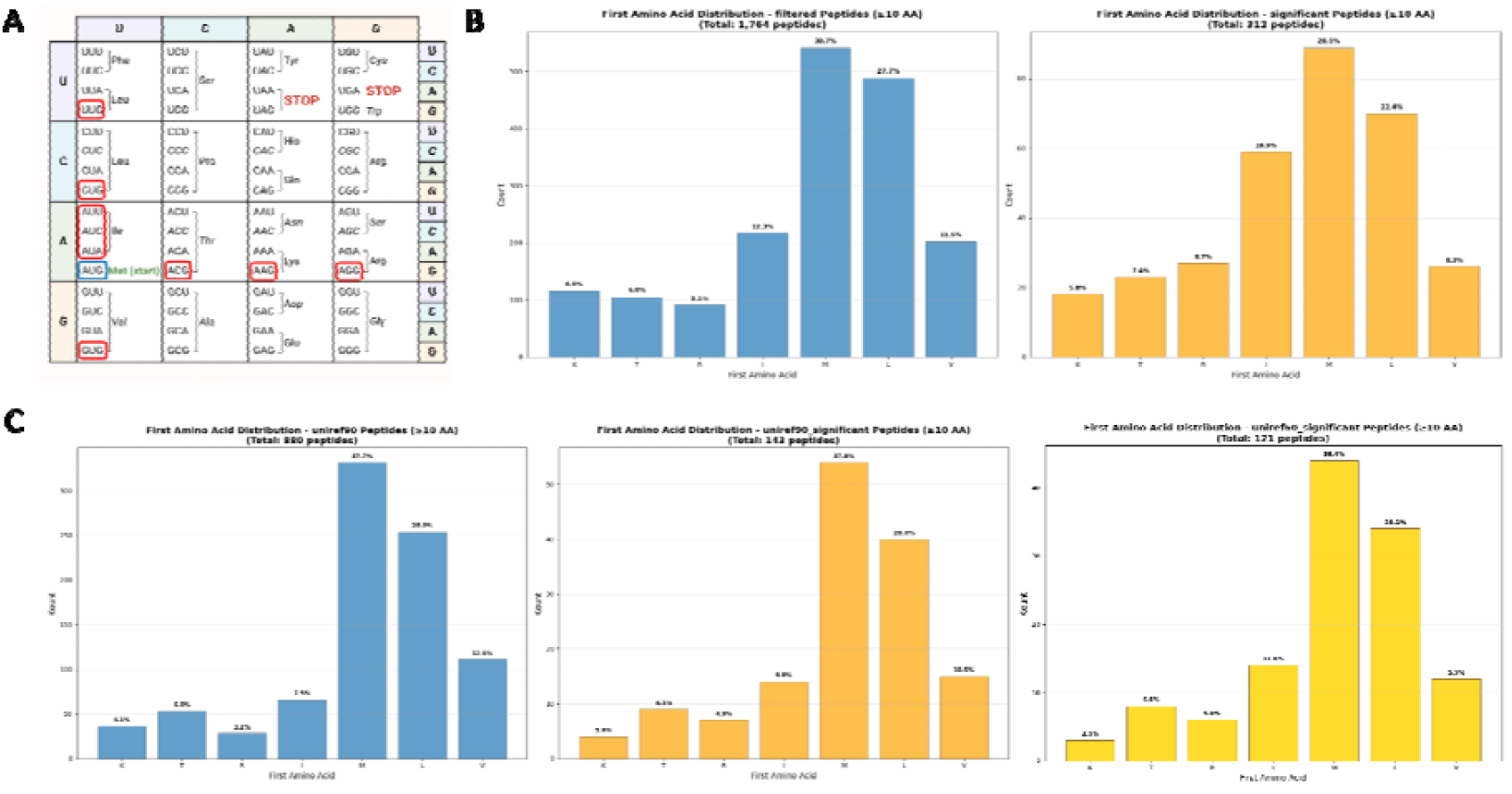
Analysis of MP initiation codon. The partition of the lncRNA-MPs by the first aa is shown for MPs >10 aa. **(A)** Codon table showing all near cognate AUG (Met) in red frames. The AUG Met is colored with a blue frame. Note that there are 2 and 3 codons that are 1-nt different from AUG for Leu and Ile, respectively. **(B)** Partition of the lncRNA-MPs (left, blue) and TR-lncRNA-MPs (orange, right). The number of peptides analyzed is indicated. **(C)** Reduced set by representatives according to Ref90 protocol (≥90% aa identity), resulting in 880 peptide representatives, while the Tr-lncRNA-MPs subset was reduced to 143 MPs. Further compression based on ≥60% identity reduction resulted in 121 peptides. Note that for all analyzed lncRNA sets, the fraction of Met was the highest, followed by Leu, which is coded by 2 near-cognate codons.

### The amino acid composition of Tr-lncRNAs-MPs differs from that of canonical proteins

The enrichment of lncRNA-MPs among Tr-lncRNAs (**Fig. 1**) and other statistics (**Fig. 2**) provides no information on the similarity or uniqueness with respect to canonical proteins. We tested whether the composition of amino acids (aa) in lncRNAs is identical to that of the human proteome. We found that the lncRNA-MPs deviate substantially from the aa composition of coding genes. Out of the 20 aa, 17 were significantly different with Arg (R), Pro (P), Cys (C) and Ser (S) being overrepresented in lncRNA-MPs relative to coding proteins, while Glu (E), Asp (D) and to a lesser extent Asn (N) and Gln (Q) show the opposite trend of being underrepresented among the lncRNA-MPs. Only the frequency of His (H), Thr (T), and Phe (F) showed no significance in their relative occurrence between the two groups (**Fig. 4A**, asterisk). Even more significant variations are observed with underrepresented Tyr (Y), a critical site for post-translational modifications (PTM) of phosphorylation. Notably, the amino acid distribution profile (based on 80.1k aa) was surprisingly similar to the profile of the unfiltered, potentially noisy MPs. Such an unfiltered collection included a much larger set with 1.33M aa (**Fig. 4A**, Supplemental **Fig. S2**). We concluded that the aa composition of the confident lncRNA-MPs is quite similar to that of the entire collection of lncRNA-MPs, even though experimental evidence for most of them is missing or below statistical significance (by SmProt score).

**Fig. 4.**
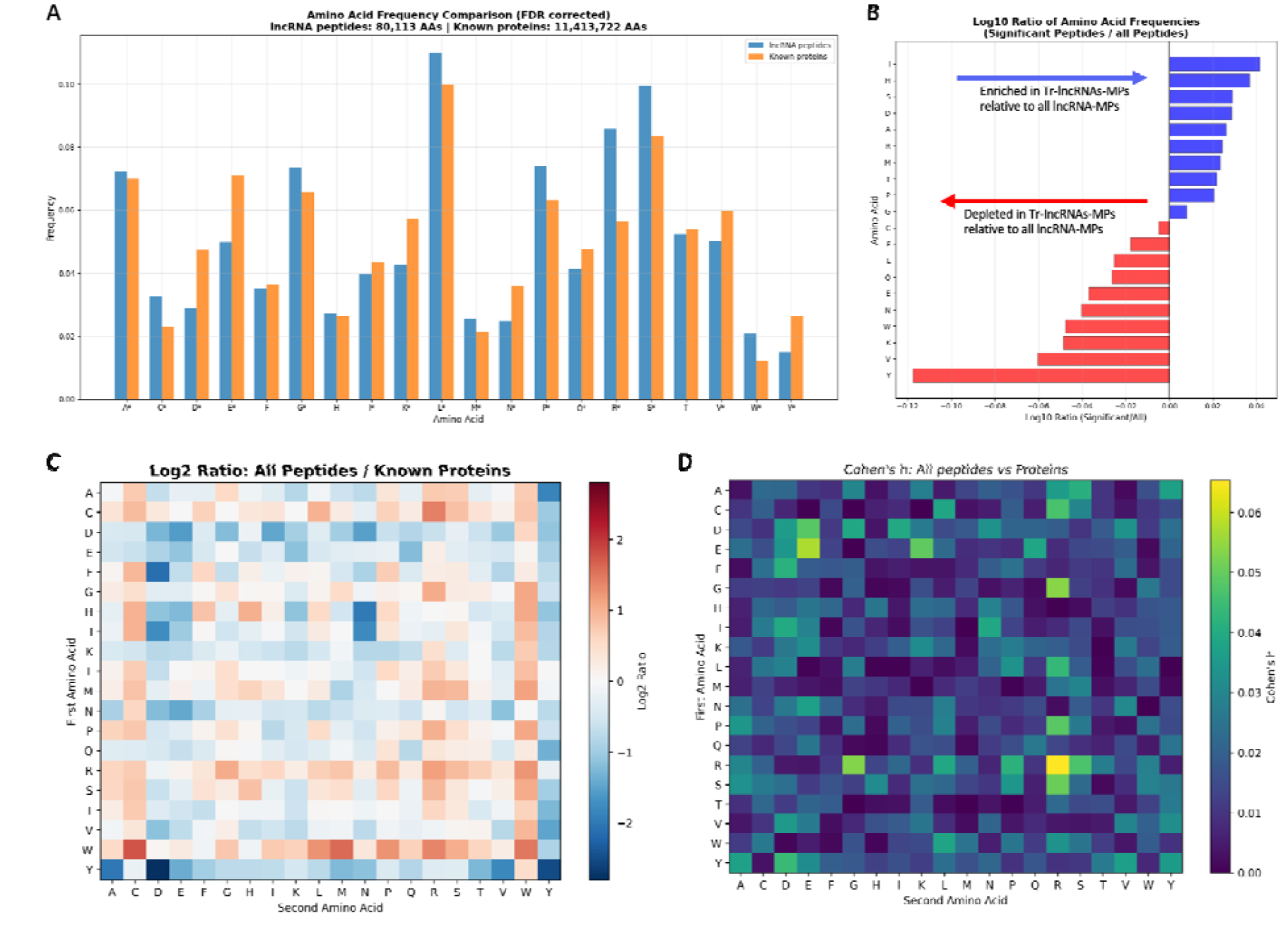
Amino acid (aa) composition. **(A)** The aa distribution of known coding proteins (orange) and high-confidence lncRNA-MPs (blue). The difference in frequency of 17 out of the 20 aa is statistically significant (marked by an asterisk). The analysis included 11.4M aa from known proteins and 80.1k aa from the lncRNA-MPs. Partitions of aa are listed in Supplementary **Table S4. (B)** The differential use of each aa in the T - lncRNA-MPs (15,230 aa) relative to all-lncRNA-MPs (80,113 aa). The relative occurrence of Ile (I) is 1.4 in the histogram, which is converted to log10(Tr-lncRNAs to all-lncRNAs ratio; x-axis). **(C)** Dipeptide occurrences in lncRNA-MPs. The number of aa for each analysis is indicated on the top of each panel. A matrix showing all 400 pairs of dipeptides and their log_2_(ratio of lncRNA-MPs and human proteome). **(D)** A matrix showing the Cohen’s h test for the deviation of the dipeptides from the human proteome (with 11.44 M aa) and the effect size. Values range from 0 to 0.07, where 0 indicates no effect and 1.0 indicates maximal effect.

Finally, we tested whether there are significant changes in the aa distributions of all lncRNAs and Tr-lncRNAs. **Fig. 4B** shows that Tr-lncRNAs are unique in their composition and significantly enriched in Ile (I), His (H), and others, while strongly depleted of Tyr (Y), the (V, K, W) among others. We concluded that the Tr-lncRNA group cannot be considered a naive sampling of lncRNA-MPs. Note that while the Tr-lncRNAs account for less than 20% of the aa amount (15.2k vs 80.1k), they exhibit strong and unique statistical characteristics. We confirmed that the results are robust and statistically valid. In Supplemental **Table S1** all confident lncRNA-MPs (1,782 peptides) and the subset of the 314 peptides belonging to Tr-lncRNAs are listed (Supplementary **Table S3**).

Despite the relatively small set of peptides that are considered as confident MPs, we tested the frequency of the dipeptides (400 values) statistics relative to the human proteome (11.41M). **Fig. 4C** shows a heatmap by log_2_-transformed frequency ratios for dipeptide sequences from all confident lncRNA-MPs relative to all protein-coding. Positive values indicate enrichment in the lncRNA-MPs, while negative values reflect the depletion relative to the entire human proteome. The strongest observation is the over-representation of the combination of Trp (W) and Cys (C), and to a lesser extent, also pairs with Arg (R). To avoid biases that reflect the default occurrence of any of the dipeptides, we transform the raw data into a statistical heatmap that captures the effect size by Cohen’s d (**Fig. 4D**). Note that the scale across all pairs of aa is quite modest (Cohen’s d <0.07), which quantifies a low difference between the two mean groups. Still, maximal values were associated with [RR], [EE], but also [GR] and [RG]. Note that despite minor differences in the dipeptide occurrences of lnc-RNAs vs. Tr-lncRNAs, the statistical difference between the Tr-lncRNA-MPs and the general list of lncRNA-MPs is negligible and failed to show statistical significance. Supplemental **Fig. S2** confirms the occurrences (by absolute numbers) of any of the 400 dipeptides from lncRNA-MPs and the dipeptide enrichment by Chi-square results of lncRNA-MPs relative to the entire human proteome.

We conclude that the pattern of dipeptides of lncRNA-MPs slightly deviates from the human proteome and that the subset of Tr-lncRNAs dominates the signal of the lncRNA-MPs. We further identified dipeptides that tend to be used in lncRNA-MPs. We claim that the predicted ORFs in lncRNAs are not merely random but exhibit sequence signatures characteristic of biologically active protein-coding regions.

### Enriched 4-mer amino acid sequence specifies lncRNAs that are signified by a large number of MPs

Based on the dipeptides statistics (**Fig. 4C**), we asked whether there are 4-mer among the confident lncRNAs and their Tr-lncRNA subset. Theoretically, there are 160k combinations of 4-mer (20^4) for all-lncRNAs that include 1.34M aa. We focused on a number of 4-mer that are unique to the high confidence MPs from the lncRNAs set.

**Fig. 5** (and Supplementary **Table S4**) shows the occurrence of 4-mers according to their uniqueness to the lncRNA-MPs as well as a few 4-mers that are abundant in this group (Supplementary **Table S5**). Despite the fact that the human proteome covers 11.41M aa and the lncRNA-MPs cover only 80.1k aa (only 0.7% relative to the human proteome), 3 cases are unique to lncRNA-MPs (IMQC, CPMF, MWNC, **Fig. 5A**, red symbol). Under a simplified estimation, the p-value to obtain 3 unique 4-mers in such a collection is 8.3e-75. There are 4 groups of 4-mers that we have considered. A set of extremely abundant 4-mers in the human proteome, which were not represented in the MPs (**Fig. 5A**, green symbols), and very abundant 4-mers in the proteome, but rare among the lncRNA-MPs (**Fig. 5A**, example KKKK, TGEK, blue background) will not be further discussed.

**Fig. 5.**
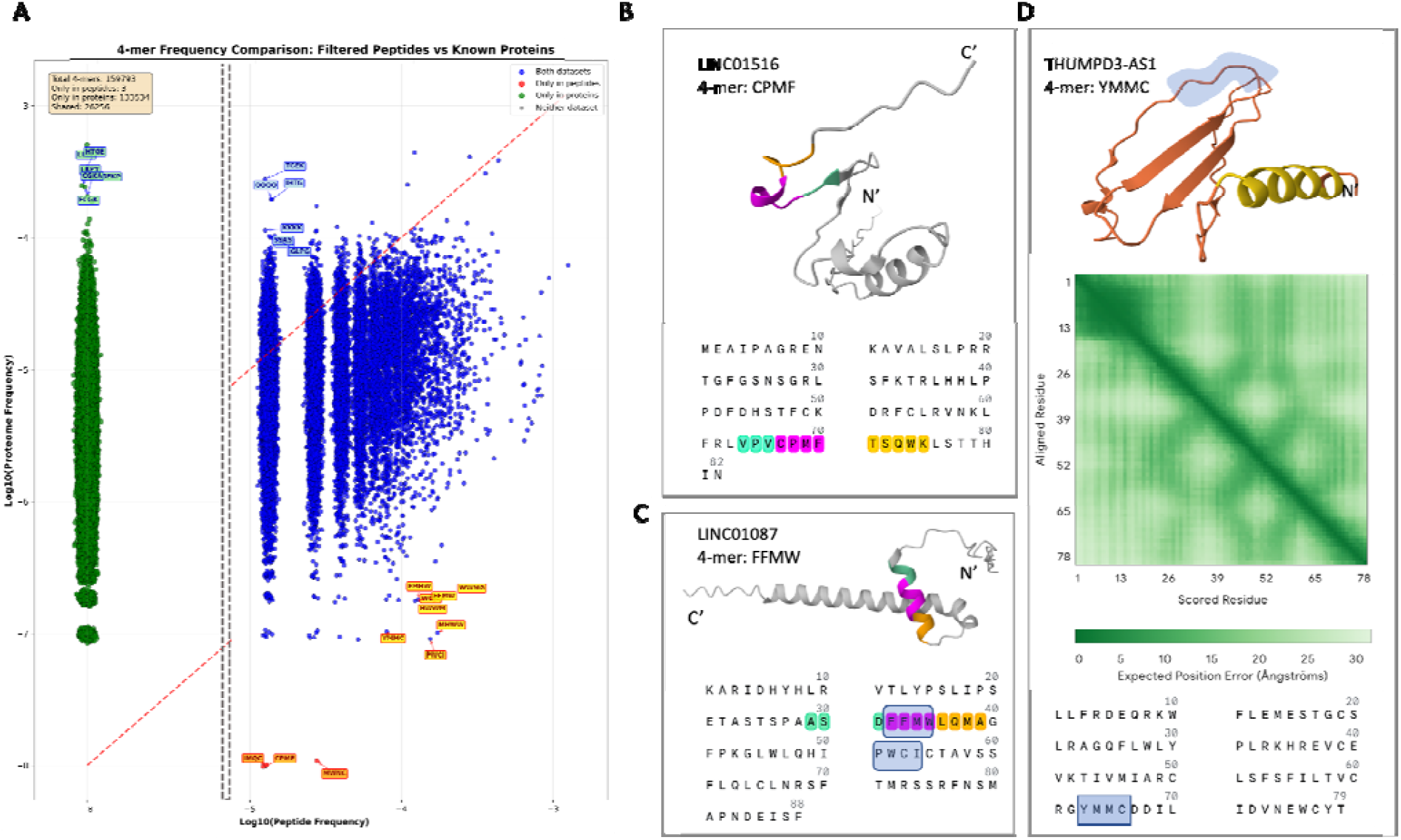
Enrichment and Structural Analysis of 4-mer sequence in lncRNA-MPs. **(A)** Scatter plot showing the frequency of 4-mer in all confident lncRNA-MPs relative to the human proteome, clustered into four colored groups. In green color are 4-mers identified exclusively in the human proteome. The sequences of the few highly abundant 4-mers are indicated. Statistical summary of the observations from the two groups (proteome and MPs) is shown. Note that the x-axis is non-continuous (separated by dashed vertical lines). **(B)** Structural prediction for lncRNAs of LINC01516 for the longest MP (82 aa) using AF2. The unique 4-mer of CPMF is colored in magenta. **(C)** Structural prediction for lncRNAs of LINC01087 for MP (88 aa) using AF2. The enriched FFMW and PWCI are marked as highly abundant in the lncRNA-MP set (shaded blue, colored in magenta. The pTM is 0.3. **(D)** Structural prediction for THUMPD3-AS1’s peptide with 79 aa. The prediction structure is of low (yellow) and very low (orange) 3D confidence according to AF2. The unique 4-mer of YMMC is shaded blue on the structure and sequence (bottom). The heatmap depicts the expected positional error (in Å) between residue pairs, where darker green indicates higher confidence. Darker colors outside of the diagonal reflect segments with local contact. The model has low-medium overall confidence (pTM = 0.24). Note that the marked 4-mer is expected to be exposed and not in contact with other segments of the peptide.

**Fig. 5B** shows the expected structure of the lncRNA-MP that includes a unique 4-mer CPMF. It was found in LINC01516, a poorly studied lncRNA with an exceptionally long transcript (8 exons, multiple transcripts). Testing its folding capacity confirmed the uncertainty in its structure (AlphaFold 2.0, AF2). There is another set of 8 sequences (4-mers) that were exceptionally abundant among the lncRNA-MPs and poorly represented in the human proteomes (**Fig. 5A**, yellow background). **Fig. 5C** shows the occurrence of FFMW in LINC01087 (88 aa, also called ERα-regulated long noncoding RNA 1). The LINC01087 is an estrogen-responsive gene, where the transcription is strongly up-regulated by ERα signaling and is strongly associated with the ER+ breast cancer subtype. The sequence is part of an α-helix, and the entire peptide is assigned with an intermediate structural confidence (pTM 0.3). **Fig. 5D** is an interesting example of a 4-mer sequence of YMMC that appears in several peptides of the THUMPD3-AS1 gene. This THUMPD3-AS1 lncRNA, also called SETD5-AS1 (genomic locus Chr3p25.3), was mentioned in the context of cancers, suggesting an oncogenic role across various cancer types^22^ . Still, all studies attributed its role to its competing endogenous RNAs (ceRNAs) function in relation to specific miRNA sponging that are involved in tumor progression.

The strict selection of confident lncRNA-MPs was essential to our analysis to secure the relevance of MPs and to better characterize their properties. Even with the small collection of 4-mer (**Fig. 5**), we observed that many of the high-occurrence sequences derived from the same lncRNA gene. For example, the LINC01087 was reported with 25 different peptides (among the unfiltered lncRNA-MP collection, the longest is of 88 aa; **Fig. 5C**). In another case of the enriched 4-mer CMTM, all 11 peptides that contain this sequence were derived from SLC9A3-AS1. This lncRNA is located at an exceptionally active genomic segment (∼10k nt) that produces many lncRNA-MPs.

### A set of lncRNAs represented by a large set of candidate peptides

Among the 1,782 filtered high-confidence MP sequences, there are 314 Tr-lncRNAs (Supplementary **Table S3**). We focused on the top listed lncRNA maximal number of MPs per gene. Inspecting the top 10 lncRNAs (**Table 1**) revealed that the majority of them are relatively long (>30 aa). They derive from a relatively long genomic region (with large variation) and transcript length (average 4,800 nt). We conclude that while there are as many as 1,782 peptides among the studied lncRNAs, the 10 lncRNAs that are enriched in MPs (**Table 1**) account for 16.7% of them. Note the extreme case of TTN-AS1 with 92% out of the overall 60 MPs, which are >30 aa (**Table 1**).

**Table 1.** Lists the top 10 Tr-lncRNA-MPs that carry a high number of MPs per gene, along with their genomic and transcriptomic features

| GeneID<br>(Ensembl) | Gene<br>Symbol | Genomic<br>size (kb) | Transcript<br>(nt) | Max<br>Length (aa) | # of MP<br>>10 aa | # of MP<br>>30 aa |
| --- | --- | --- | --- | --- | --- | --- |
| ENSG00000237298.10 | TTN-AS1 | 258 | 6,297 | 98 | 60 | 55 |
| ENSG00000226688.7 | ENTPD1-AS1 | 357 | 5,112 | 99 | 35 | 22 |
| ENSG00000245694.10 | CRNDE | 86 | 6,325 | 75 | 35 | 29 |
| ENSG00000247828.8 | TMEM161B-AS1 | 170 | 15,798 | 74 | 28 | 12 |
| ENSG00000249859.11 | PVT1 | 393 | 2,149 | 91 | 27 | 18 |
| ENSG00000196756.13 | SNHG17 | 15 | 2,261 | 83 | 22 | 17 |
| ENSG00000257621.8 | PSMA3-AS1 | 160 | 2,475 | 84 | 22 | 19 |
| ENSG00000242125.3 | SNHG3 | 12 | 2,614 | 60 | 20 | 12 |
| ENSG00000255717.7 | SNHG1 | 4 | 2,812 | 66 | 20 | 12 |
| ENSG00000233461.6 | AL445524.1 | 7.5 | 2,172 | 83 | 19 | 18 |

It is important to note that several of the listed genes from **Table 1** were studied in the context of lncRNA roles in cancer identity (e.g., PVT1), immunomodulation (e.g., ENTPD1-AS1), proliferation (e.g., TTN-AS1), and more. TTN-AS1 is represented by 60 individual peptides (**Table 1**), and it is a major Tr-lncRNA signifying HNSC E→L transition. We demonstrate the richness of lncRNA-MPs that belong to the SNHG family members (3 genes). We identified in the top MP-enriched genes 3 members of the nucleolar RNA host genes (SNHG3, SNHG17, and SNHG1). These genes act as key drivers of cancer progression. SNHG3 is upregulated mostly in breast, liver, and lung, and SNHG17 in gastric, colorectal, and lung. The expressed members mainly function as competing endogenous RNAs (ceRNAs) that promote epithelial–mesenchymal transition (EMT), proliferation, and immune evasion. SNHG1 forms a regulatory network confirming its oncogenic function in colorectal cancer and other cancer types. **Fig. 6A** shows the structural prediction of SNHG1 (65 aa). It shows that the long alpha helix is of very high confidence (colored blue) and the N’-terminal is somewhat less evident. Using SwissModel, there was a single model that indicated a well-organized tetrameric packed structure (**Fig. 6B**). Inspecting the model revealed the expected protein to be a membrane protein that is composed of 4 main chains in a repeated coordinated structure resembling the human TRPM4^23^. The predicted tetramer was solved by cryo-EM and validated as a cation channel crossing the membrane (**Fig. 6C**). TRPM4 is a calcium-activated, nonselective monovalent cation channel that acts in membrane potenti l and calcium signaling in diverse systems. We concluded that the lncRNA-MPs of SNHG1 may interfere with the function of the unaltered channel by interfering with the oligomeric structure of TRPM4^23^.

**Fig. 6.**
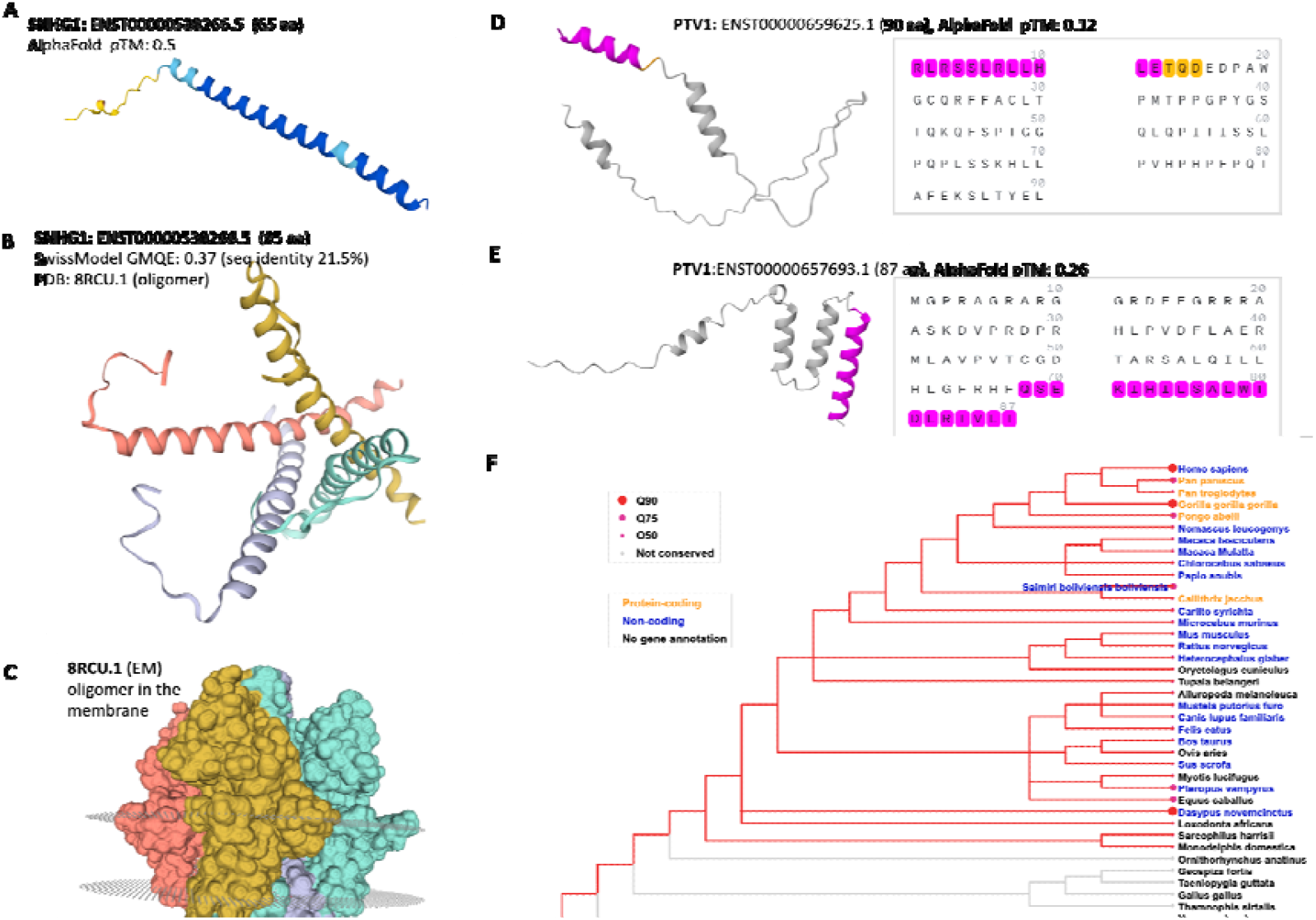
Structure and phylogenetic analysis of lncRNA with a rich collection of MPs. **(A)** Structural prediction for the SNHG1 sequence (65 aa) by AlphaFold 2.0. The structure prediction displays a very long alpha-helix (high confidence, pTM score of 0.5). **(B)** SwissModel builds a model for a tetramer structure (as in A) that is positioned to form a conducting channel. **(C)** The full model complex of the human TRPM4 was identified PDB: 8RCU.1 as a template (note the low sequence identity). The structure was based on the cryo-electron microscopy (cryo-EM) structure of the TRPM4 oligomer embedded in a membrane. TRPM4 is a calcium-activated monovalent cation channel involved in cellular excitability and signal transduction. **(D)** Sequence of transcript variant PVT1 (90 aa). The prediction is of low confidence (pTM =0.12). **(E)** Sequence of transcript variant PVT1 (87 aa). The prediction is of low-medium confidence (pTM =0.26). Note that there is no overlap in sequence between the two transcripts of PVT1. Helical regions shown in magenta match the predicted structure by AF2. (F) Phylogenetic tree illustrating the evolutionary conservation of PVT1 homologs across species. Species annotated by homologs that are protein-coding are highlighted in orange. The tree reveals strong conservation among mammals (with some exceptions). Still, the lncRNAs in humans are homologs to coding genes in primates.

All genes listed in **Table 1**, including SNHG1 members, were not implicated to act by their potential proteins. An exception is the PVT1 encoded peptide that was recognized as a tumor antigen in colon cancer ^24^. Remarkably, PVT1 spans an extremely long genomic region (**Table 1**) and its transcript is exceptionally long (total 13 exons, some transcripts reach 19-20k nucleotides). Several MPs that are derived from PVT1 do not overlap in sequence. These MPs are encoded from different exons and the alternative use of open reading frames. Two of the long overlapping MPs of PVT1 are illustrated in **Fig. 6**. The prediction from AF2, shown in **Fig. 6A** and **Fig. 6B** confirms that both sequences are poorly structured (belong to clusters 1 and 2, respectively; Supplementary **Fig. S4,** Supplementary **Table S7.**). The predicted sequences are unpacked with low to modest quality for folding accuracy. **Fig. 6C** shows the phylogenetic tree of the PVT1 gene. It is interesting to note that some primates are annotated as coding genes, while in humans, gibbons, macaque, and other Old World monkeys are marked as lncRNAs. Interestingly, in Callithrix, a genus of New World monkeys, PVT1 was annotated as a coding gene. Homologs were not found in treeshrews (a small mammal), rabbits, and a few other mammals, suggesting high diversity of PVT1 (Supplementary **Table S3**).

**Fig. 7B** presents the AlphaFold2-predicted structure of TTN-AS1, a notable Tr-lncRNA implicated in the epithelial-to-luminal (E→L) transition of HNSC. The model predicts multiple β-strands with substantial inter-strand contacts, as well as a long, intrinsically unstructured tail segment. **Fig. 7C** depicts the estimated positional error between all residue pairs. Overall, the model exhibits low confidence in residue placement (pTM = 0.14), indicating limited reliability of the predicted segment positions. The accompanying heatmap illustrates the expected pairwise positional error (Å) between residues, with darker green tones beyond the diagonal representing greater uncertainty.

**Fig. 7.**
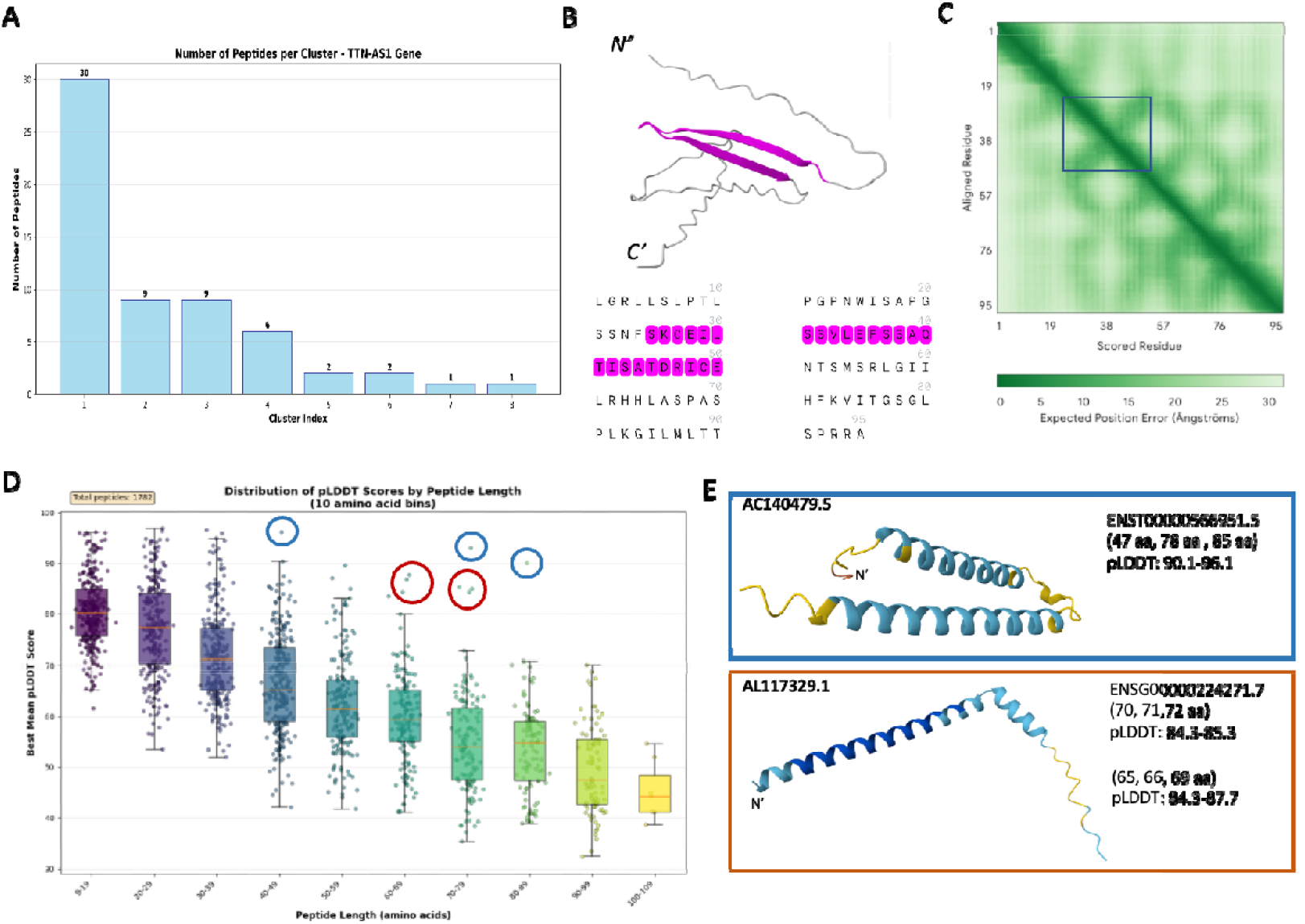
Structural prediction analysis. **(A)** Clustering by sequence identity (>60%) of TTN-AS1 Tr-lncRNA. 8 clusters account for 60 MPs. **(B)** Structural prediction for Tr-lncRNAs of TTN-AS1 for the longest MP (from cluster 1 in A) using AlphaFold 2.0. TTN-AS1 is represented by 60 individual peptides **(Table 1)**. Sequence of the 95 aa MP of TTN-AS1 colored (**β**-sheets shown above). The TTN-AS1 is a Tr-lncRNA that is specialized in HNSC E→L transition. **(C)** A heatmap depicts the expected positional error (in Å) between residue pairs, where darker green indicates higher confidence. Darker colors outside of the diagonal reflect segments with local contact. The lighter regions suggest greater uncertainty. The model has low overall confidence (pTM = 0.14).**(D)** Box plot for the pLDDT from AlphaFold2 (AF2) by MP length bins. The longer the MPs, the lower the mean pLDDT is (on average). The red circles mark 6 MPs whose length range between 60-79 aa and are indicated with high pLDDT, assigned to lncRNA AL117329.1, and the blue circles are associated with AC140479.5 with outstanding high pLDDT (0.91). **(E)** The AF2 prediction of AC140479.5 (85 aa, blue circles in D) and AL117329.1 (69 aa, red circles in D). The representative sequences of the marked clusters of MPs are signified by relatively long α−helices. The bluish color of the 3-D structure indicates a prediction of high confidence pLDDT.

We evaluated whether pLDDT, AlphaFold’s per-residue confidence score, could serve as an indicator of the predicted MPs’ three-dimensional structure. To this end, we analyzed the correlation between pLDDT values and peptide length (**Fig. 7D**). Supplementary **Table S1** lists all 1,782 peptides along with their AF2-derived statistical metrics. An inverse relationship emerged: longer MPs tended to exhibit lower pLDDT scores. We further examined MP instances with exceptionally high pLDDT values and found that such sequences were enriched in α-helical structures. Notable high-pLDDT outliers include AC140479.5 (85 amino acids; blue circles in **Fig. 7D**) and AL117329.1 (69 amino acids; red circles in **Fig. 7D**). Both lncRNAs remain poorly characterized; however, differential expression of AC140479.5 has been reported in FOXA2 knockouts of iPSC-derived pancreatic cells, suggesting a potential link to cell identity and organ development.

We conclude that several lncRNA-derived MPs are foldable PMs. Consequently, their protein products may serve as protein-based signatures in specific cancer contexts and could participate in promoting or modulating protein–protein interactions (PPIs).

### Predicted signal peptide sequence highlights MPs of the secretory system

We applied SignalP 6.0 to predict signal peptides (SPs, see Methods) for the 1,782 high-confidence MPs. Only 6 MPs were positively predicted to have a signal peptide with high probability, accounting for 4 unique lncRNA genes: ZNF433-AS1, AC020928.1, LOXL1-AS1, and SNHG17 (Supplementary **Table S1,** Supplementary **Fig. S3**). A clear cleavage site in all 4 proteins peaks around residues 20–30 aa, indicating that these MPs could be secreted or targeted to the endoplasmic reticulum (ER). LOXL1-AS1 is an antisense of LOXL1 that plays a role in extracellular matrix remodeling and fibrosis (by crosslinking collagen and elastin). ZNF433-AS1 is poorly studied but potentially affects the expression of ZNF433, which is involved in chromatin modelling. The SNHG17, like other SNHG members, is implicated in cancer development and progression. Collectively, we show that only a negligible fraction of lncRNA-MPs exhibits classical SP features.

### Overlooked Tr-lncRNA-MP shows similarity to known functional protein domains

In our analysis, we determined strict thresholds for MPs and Tr-lncRNAs. Altogether, we discuss a selected confident set of 1,782 high-confidence MPs, among them only 314 peptides (72 genes) matched the Tr-lncRNA definition. We tested the possibility of identifying MPs that can be functionally interpreted. To this end, we applied the Swiss Model that created the most likely 3D models, as well as AF2, which provides an estimate for the foldability of the query sequence, as well as an error model for each pair of aa in the sequence.

**Fig. 8** highlights three genes represented by multiple peptides originating from Tr-lncRNAs, integrating 3D modeling using AF2 predictions with SwissModel-derived best templates and associated confidence statistics (GMQE, a normalized metric comparing the query to its best template). **Fig. 8A** illustrates a ubiquitin-like structure supported by extremely high sequence similarity and elevated GMQE scores (**Fig. 8B**). Notably, CDKN2B-AS1 resides on the reverse strand of CDKN2B, which encodes p15INK4b, a well-characterized cyclin-dependent kinase inhibitor. p15INK4b functions through inhibition of CDK4 and CDK6, the principal regulators of the G1→S phase transition of the cell cycle. **Fig. 8C** depicts an understudied MP folded into a compact four-helical bundle, one helix of which (**Fig. 8C**) shows strong homology among primates for an uncharacterized ∼40-residue domain (positions 45–85, **Fig. 8D**). **Fig. 8E** presents a predicted 3D structure resembling classical RNase H (found in bacteria, viruses, and other organisms). Collectively, these findings suggest that MPs functionally linked to cancer progression or metastatic transition may act by mimicking known small proteins (e.g., RNase H, ubiquitin), thereby potentially interfering with or promoting protein– protein interactions during cancer progression.

**Fig. 8.**
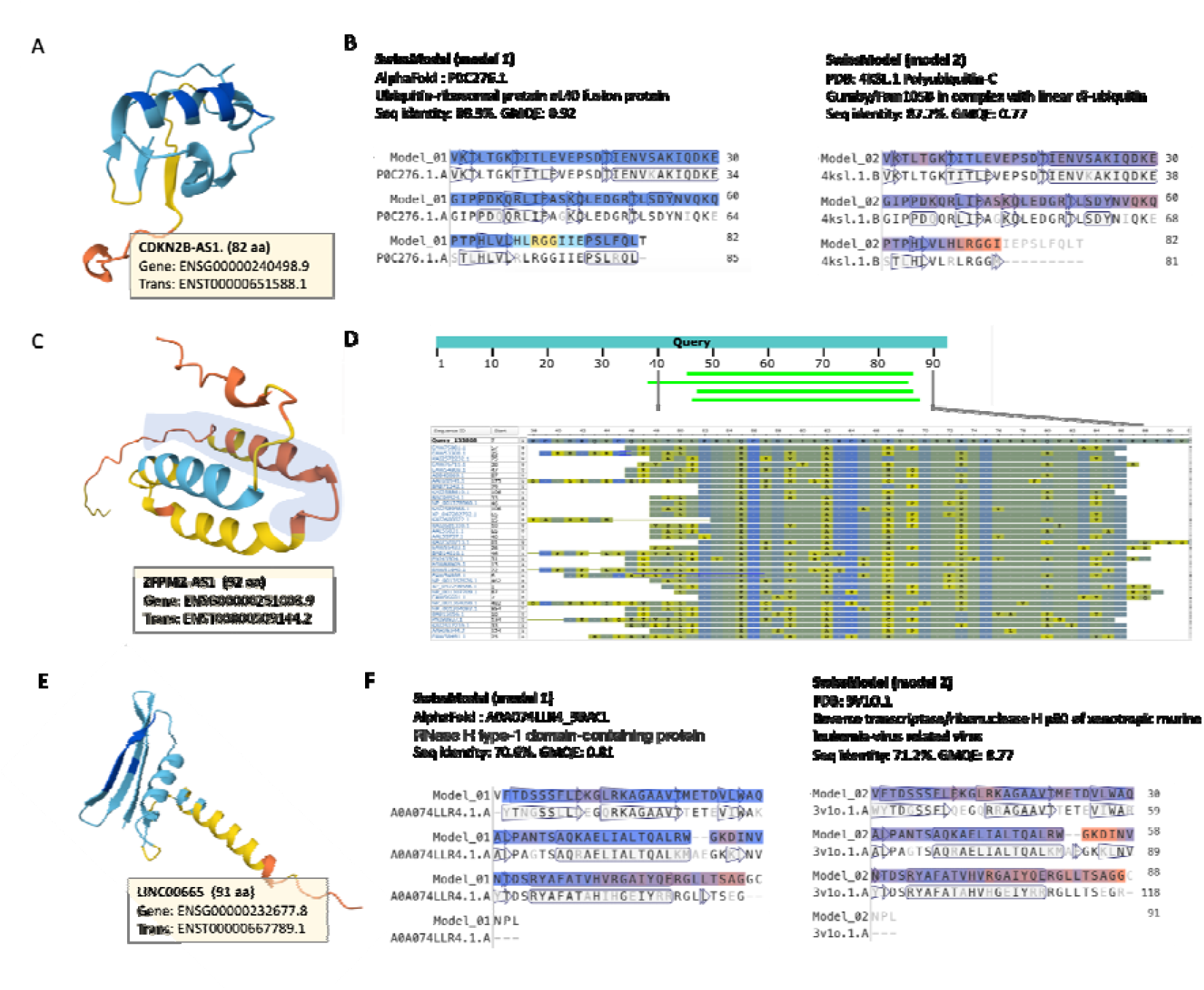
Predicted micropeptides encoded by selected Tr-lncRNAs **(A)** CDKN2B-AS1 (ENSG00000240498.9; transcript ENST0000051588.1; 82 aa). The predicted 3D fold shows mixed α-helical and β-strand elements. **(B)** A predicted peptide with its best structural homologs according to SwissModel, the two best models of ubiquitin. The conserved residues are indicated by a blue shading, and regions with low similarity are colored red. **(C)** ZFPM2-AS1 (ENSG00000251003.9; transcript ENST0000059144.2; 92 aa). The predicted structure is composed of a few well-packed alpha-helices of unknown function. The shaded helix strongly resembles a short domain of 40 aa, marked by light green along the position of the sequence. **(D)** Multiple sequence alignments (MSA) of the overlapping helical domain. Blue color indicates that the query and the listed homologues share identical aa. The yellow color marks a consensus among the homologs (not identical to the query sequence). The high degree of sequence similarity spans proteins from humans and other primates.; **(E)** LINC00665 (ENSG00000252677.8; transcript ENST0000067789.1; 91 aa). The predicted models from SwissProt show the overlap in sequence and structure with RNase H type-1–like fold; **(F)** Pairwise alignment with the top two models built by SwissModel. The similarity to RNase H type-1 domain-containing protein and reverse transcriptase/RNAse H is evident. The conserved aa are colored blue, and the high diversity sequences are colored red.

### Pan-cancer Tr-lncRNAs are enriched with MPs

Pan-cancer analyses reveal Tr-lncRNAs shared across multiple cancer types and cellular transition^21^ . We investigated whether MPs derived from these top pan-cancer Tr-lncRNAs might contribute to cancer progression. **Fig. 9** presents a Venn diagram of the top 50 Tr-lncRNAs identified across four T-stage transitions and three M-state transitions in 17 cancer types, regardless of MP presence. We then assessed whether MPs were enriched within these top pan-cancer gene sets. The Venn diagram (**Fig. 9A**) illustrates the overlap between the T-stage and M-state pan-cancer gene groups (29 shared genes; Supplemental **Table S6**), representing a highly significant enrichment (hypergeometric test, p-value <1e-30). Although the numbers were smaller and did not reach statistical significance, we observed that 5 of the 72 Tr-lncRNAs encoding high-confidence MPs overlapped with the pan-cancer lists (LINC01234, HAND-AS1, XIST, UCA1, and HOXA11-AS)

**Fig. 9.**
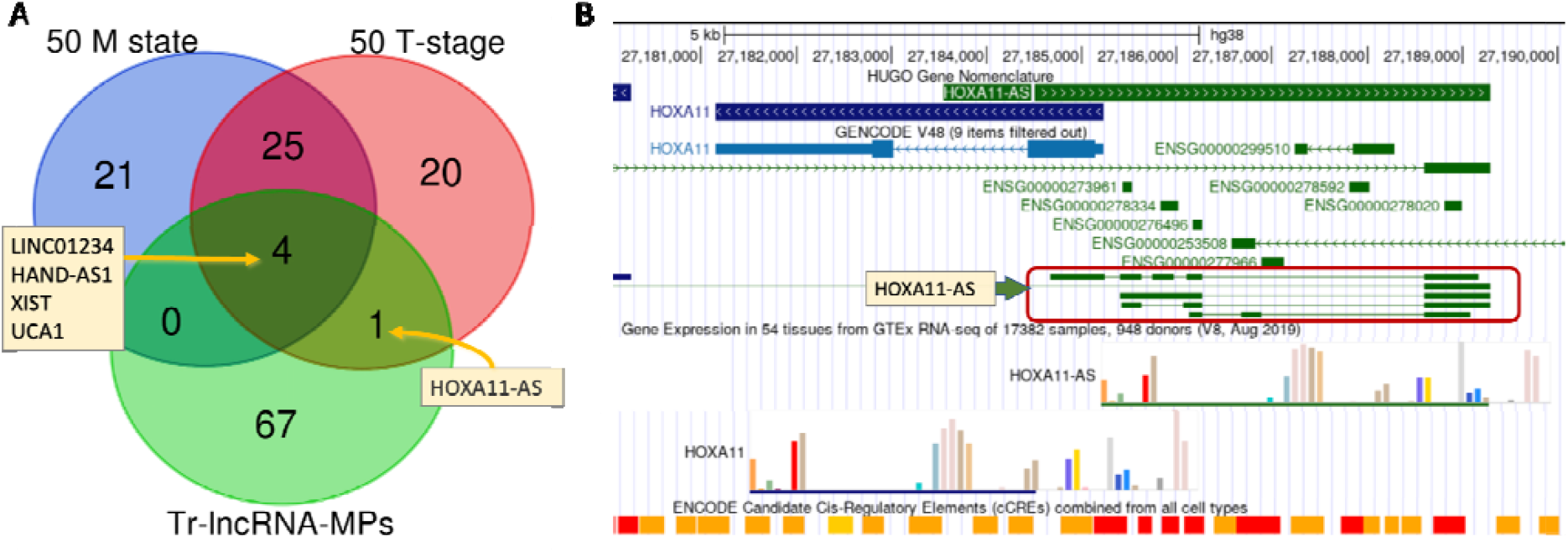
Pan cancer analysis of Tr-lncRNAs and MPs of >30 aa. **(A)** The Venn diagram emphasizes the shared and unique Tr-lncRNAs assigned as pancancer. The top Tr-lncRNAs associated with M-state and T-stage transitions are compared (top 50 Tr-lncRNAs for each group). The 72 genes that are represented by 314 MPs >10 aa are shown, and the overlapping set of Tr-lnRNAs names are listed. **(B)** UCSC genome browser view of HOXA11 and HOXA11-AS. The tracks cover 9,000 nt from Chr7. The HUGO track shows the segment of overlap of the two genes. GENCODE V48 shows the multiple transcripts of HOXA11-AS (green color). Gene expression across 54 tissues from GTEx is shown. Each colored bar is a different tissue. A strong coherence of HOXA11 and HOXA11-AS is observed. The ENCODE track is colored according to the regulatory element annotations with red, orange, and yellow blocks that mark promoter-like signature (PLS), proximal enhancer-like signature (pELS), and distal enhancer-like signature (dELS), respectively.

**Fig. 9B** shows the dense region of the HOXA11 and the HOXA11-AS. There are several observations from the genomic perspective: (i) HOXA11-AS is expressed with several versions and an alternative spliced antisense transcript (red frame, all 5 transcripts were listed as highly confident MPs). (ii) HOXA11 and the HOXA11-AS overlap in their transcripts on the first exon that includes the Met start codon of HOXA11. (iii) This region is transcriptionally regulated and includes a cluster of promoters (**Fig. 9B,** ENCODE track, red) and enhancer/promoters (ENCODE track, orange) segments. (iv) The expression profiles of the HOXA11 and the HOXA11-AS (shown by the GTEx track) are extremely similar, which suggests their coupling. Specifically, higher expression is detected in the cervix, colon, thyroid, artery-tibial, bladder, and others. We concluded that the transcriptional coordination, as determined by the GTEx track, extends to the MPs encoded by the HOXA11-AS. Among the other genes (LINC01234, HAND-AS1, XIST, and UCA1), only LINC01234 was suggested to encode a protein (called MBOP) that promotes colorectal cancer via MAPK signaling. The protein capacity of the other genes was not discussed, and their relevant function was exclusively attributed to their transcriptional levels and impact on signaling in cancer ^6^.

## Discussion

This study provides a comprehensive view of the emerging role of lncRNA-encoded micropeptides (lncRNA-MPs) and reveals their complex expression landscape across cancer progression. Although traditionally classified as noncoding, an increasing body of evidence indicates that many lncRNAs encode functional peptides ^25^. Here, we focus on high-confidence lncRNA-MPs, which represent only a small fraction (∼10%) of the unfiltered pool of MP candidates from the lncRNA transcripts. Using TCGA data ^21^, we identified 2,399 genes termed Tr-lncRNAs that are unfiltered genes. These genes’ transcripts were markedly enriched for unfiltered predicted MPs (60%) compared with all lncRNAs (21%; Fig. 1A). There are 2,935 peptides that are high-confidence (from an initial collection of 19,920), we suggest these may expand the unexausted cancer-associated MP candidates. Notably, many lncRNA-derived peptides may have been overlooked because of noncanonical features such as alternative start codons (present in >50% of cases; Fig. 3). Additional experimental approaches, such as mass spectrometry and immunosurveillance, can surface epitopes and may uncover further MPs of interest ^26^.

Many lncRNAs were associated with multiple peptides (Table 1). To reduce redundancy, we applied Ref90 and Ref60 clustering (minimum peptide size ≥10 aa), thereby improving the signal-to-noise ratio of high-confidence MPs (Fig. 3). Integrating TCGA data across 17 cancer types yielded 478 lncRNAs encoding 1,782 peptides. Sequence-based clustering, AF2, and Swiss-Model predictions helped define a “hidden peptidome.” The strong correlation between peptide size and pLDDT scores (Fig. 6D) enabled the identification of peptides whose AF2 pLDDT values deviated from the mean (Supplementary Table S1). These peptides typically form robust α-helical structures (Fig. 5E), suggesting increased stability and potential for protein–protein interactions.

The average length of the lncRNA-MPs in this study was 45 aa (Supplementary Table S1), making functional annotation particularly challenging. Longer peptides (>80 aa) were more suitable for homology searches and structural inference (Fig. 7). There are sporadic examples in the literature for shorter functional lncRNA-derived MPs. For example, the 48 aa TUNAR peptide, implicated in neuronal differentiation and ER calcium homeostasis in pancreatic β cells ^27, 28^. In another case, a 17 aa peptide from MIR155HG was shown to suppress autoimmune inflammation ^29^. A sort peptide (21 aa) from MLLT4-AS1 was associated with survival in breast cancer ^30^. These examples illustrate that even very short MPs can exert important biological effects, but ranking them will require new approaches to decipher their functions. While most known MPs affect intracellular signaling, those containing canonical signal peptides may also be secreted via extracellular vesicles (EVs), influencing the tumor microenvironment. Such MPs could serve as attractive diagnostic or prognostic biomarkers. Importantly, peptides lacking canonical signal peptides may still be secreted via EVs 31.

Several lncRNAs have been reported in the context of cancer ^32^ and as unexplored potential therapeutic targets. Our study highlights a subset of cancer-associated lncRNAs, including members of the SNHG family. Although SNHG genes are strongly implicated in cancer biology, their MP potential has been largely unexplored. We identified a rich collection of MPs within SNHG3 (20 peptides), SNHG17 (22 peptides), and SNHG1 (20 peptides). Notably, SNHG17 harbors a putative signal peptide that may facilitate secretion (Supplemental Fig. S3). SNHG genes are recognized oncogenic drivers across diverse tumor types, with SNHG17 in gastric, colorectal, and lung cancer ^33^, SNHG3 in breast, liver, and lung cancers, and SNHG1 across multiple malignancies ^34^. Overall, 7.2% of all high-confidence lncRNA-MPs originated from 13 different SNHG genes.

Several previously validated lncRNA-derived peptides, such as SPAR, HOXB-AS3, and longer MPs (BXW7-185 aa, SHPRH-146 aa) carry tumor-suppressive potential ^35^. Although our analysis focused on MPs ≤100 aa, HOXB-AS3 emerged as a strong candidate among Tr-lncRNA-MPs. We also identified LINC01234 as a pan-cancer lncRNA with high-confidence MPs, including 8 peptide variants. Recently, the LINC01234-encoded peptide MBOP (85 aa) was shown to drive colon cancer cell migration and proliferation by activating MAPK signaling via MEK1 binding and proteasomal stabilization ^36^. We anticipate that many lncRNAs encode MPs bearing structural motifs (e.g., ubiquitin, RNase H), secretion signals, and protein interaction domains, enabling them to mimic, modulate, or compete with established protein networks. These peptide functions may complement the canonical RNA-based activities of lncRNAs, such as chromatin organization, protein scaffolding, and miRNA sponging, thereby reinforcing oncogenic effects.

Discovering antigen repertoire along cancer progression is of growing interest, particularly for advancing immunotherapy. The oncogenic lncRNA PVT1 encodes a tumor antigen that elicits T-cell surveillance ^24^. Structural prediction from the Tr-lnc-MPs identified 27 MPs that failed to produce a clear folded structure. Instead, the 3D prediction tools (e.g., SwissModel, AF2) revealed several low-confidence helices and overall unpacked structures (Fig. 6D). Further support for the protein capacity of PVT1 is based on the conservation view (Fig. 6F). From a clinical perspective, high PVT1 expression correlates with poor survival in many tumor types, and PVT1 overexpression is linked to chemoresistance via apoptosis inhibition and altered signaling pathways ^37^. We propose that highly expressed lncRNA-MPs in TCGA cancer samples may enrich the repertoire of tumor antigens. Once presented on MHC class I molecules, these peptides become visible to the immune system, providing a novel layer of tumor antigen presentation ^38^.

Finally, this study underscores the limitations of current annotations. Many lncRNAs may actually represent misclassified protein-coding genes, especially in cancer-specific contexts. The definition of lncRNAs remains fluid, and pseudogenes or unspliced transcripts may infiltrate unfiltered datasets ^1^. We therefore restricted our analysis to reliable lncRNA-MPs from the SmProt dataset ^39^. Resources such as lncBook offer complementary approaches to improve MP reliability. Another limitation in functional inference of lncRNA-MP arises from the difficulty in predicting function by any structural prediction approach for peptides that are too short (e.g., <60 aa). Finally, although mass spectrometry provides the most direct evidence for MPs, identification remains limited due to low abundance, minimal peptide counts, post-translational modification, and the high level of specificity of lncRNAs expression that often match specific stages in cancer development.

## Conclusions

We confirmed that lncRNA-encoded microproteins (MPs) represent a potentially widespread layer of regulation within cancer transcriptomes. The framework developed here establishes a foundation for integrating novel lncRNA-derived proteins into our understanding of the cancer proteome. By examining expression patterns across multiple calcer transitional stages, rather than focusing solely on tumor profiles, we identified a subset of Tr-lncRNA-MPs most likely to drive cancer progression and the shift toward metastasis. Among the 314 informative MPs, we proposed putative cellular roles for tens of MPs, including ubiquitin-like activity, RNase H-related structures, and additional diverse foldable domains. We argue that the encoded peptides may influence antigen presentation, cell-cell communication, and protein–protein interactions. Extensive experimental validation and functional assays are imperative for defining and substantiating the therapeutic potential of MPs in cancer.

## Methods

### Expression and clinical data for major cancer samples

RNA sequencing (RNA-seq) data and the associated clinical annotations were obtained from The Cancer Genome Atlas (TCGA) via the UCSC Xena data portal ^19^. The RNA-seq data were generated using the HiSeq platform, employing a poly(A)+ selection protocol and paired-end sequencing strategy. Raw read counts for annotated lncRNAs were extracted and processed, and went through the TOIL pipeline, which included library depth normalization ^40^. In TCGA, the reported measures are provided as RSEM (RNA-seq by expectation maximization) for estimating gene and isoform expression levels from the raw data. The normalization is based on log2[expected_counts+1] to avoid biases from exceptionally small numbers.

Clinical metadata, including cancer type, tumor stage, and survival outcomes, were curated from the Pan-Cancer Atlas of TCGA, ensuring consistent stage classification across studies. We used the routine clinical TNM categories ^41^. We consider the T-(tumor) state and M-(metastasis) state. For the T-stages, we consider stages I to IV as the stages mark greater local invasion (and include T2a and T2b as stage II, when available). We ignored samples marked as T0 (normal, no evidence of a primary tumor) or Tis (carcinoma in situ). The M0 covers tumors that have not spread, while M1 indicates the presence of distant metastasis. We do not address the involvement of regional lymph nodes (N) due to a lack of robust classification.

### Differentially expressed transitional lncRNAs (Tr-lncRNAs)

Two types of labeling were used for the clinical annotations. The T and M system. For the T-stage, we applied each sample to its stage (I-IV), and for the M (metastasis) state, we considered M0 as the primary tumor and M1 as metastasis. We first filtered the lncRNAs to include only lncRNAs that have an expected count value of 1.0 in at least 30% percent of the samples (in at least one of the two stages of the transition) to avoid small fractional expression values likely to reflect noise. Then, we used z-scores over the log2(FC) of the means over the log(x+1) expression values of the lncRNA across samples in the two consecutive stages participating in the transition (e.g., the difference of stage T-II to stage T-I). We confirmed that the log2(FC) approximates a normal distribution. We defined lncRNA as Tr-lncRNA if it was expressed in any of the major cancer types, and the statistics of the differentially expressed gene are with a z-value >|3| for any consecutive stages ^21^. The calculation of the z-value is based on considering the mean and standard deviation of the differences of any specific cancer. The subset of the significant Tr-lncRNAs compiled from 2,399 lncRNA genes covers 7 transitions and 17 major cancer types.

### Database of micropeptides (MPs)

The unfiltered comprehensive catalog of lncRNAs was reported from GENCODE V.49 ^1^. For harmonization of annotations and unified naming, GeneCaRNA database ^42^ was used. Altogether, the GeneCaRNA reported on ∼220,000 unique ncRNA identifiers, and a gene-centric view report on >22k genes. As many as 20% of these genes are associated with MPs as determined by lncBOOK 2.0 ^43^ via GeneCaRNA catalog ^42^. Revisiting the unfiltered collection of lncRNAs was needed to maintain only the highly confident MPs for downstream analysis.

Small peptides and proteins, also called small ORF-encoded peptides (sORFs) or small encoded peptides (SEPs), were obtained from the SmProt dataset ^39^, which specializes in small proteins supported by Ribo-seq data across human tissues and cell lines. The dataset initially contained 19,920 different MPs whose peptide length is <100 aa, originating in 2,935 lncRNA genes. These MPs were filtered to include only MPs whose translation initiation site (TIS) p-value and Ribo p-value (ribosome footprint enrichment and periodicity) are <0.05. We also reduced redundancy by removing duplicates originating from multiple transcripts of the same lncRNA gene. Altogether, we report on 1,782 peptides. Among them, there are 314 peptides assigned to Tr-lncRNAs.

### SignalP 6.0

The prediction of the signal peptide sequence relies on the properties of the signal peptides (SPs) ^44^. SignalP predictor combines sequence-based features from the N’-terminal. SignalP provides a likelihood score based on 1-5 positively charged residues (Arg and Lys), hydrophobic core (with tracks of Leu, Ile, Val, and Phe), and the position of a cleavage site (CS), which includes small, neutral residues for SP peptidase. Specifically, the recognition signature for signal peptidase I is based on the presence of Ala, Ser, or Gly at -1 and -3 positions relative to the cleavage site.

### Conservation labeling by lncBook

The lncRNA homologous sequences are based on the alignment length (≥50 nt) and a minimal length of >20% of the lncRNA transcripts. For a specific species, the homologous sequences are determined relative to the Q50 threshold for introns. A sequence in another species was called homologous if its alignment length and identity exceeded the intronic Q50 threshold (the median level of intron conservation) ^43^. For conservation of Q75, the lncRNA (exon/intron) has a conservation score higher than at least 75% of comparable to the bulk of lncRNAs. LncBook reports on ∼140k homologous genes for 22,347 human lncRNA genes. The phylogenetic tree covers 40 species from zebrafish to human.

### UniRef-based sequence clustering

To reduce redundancy among predicted peptides, we employed CD-HIT (version 4.8.1) to generate UniRef-style non-redundant sets. Input amino acid sequences were clustered at two sequence identity thresholds corresponding to UniRef definitions: 90% (UniRef90) and 60% (UniRef60). Word size was set according to CD-HIT recommendations for each threshold.

### External analysis and bioinformatics

For structural prediction, we have used AlphaFold 2.0 and the predicted Local Distance Difference Test (pLDDT) ^45^. We also applied the SwissModel server for functional relatedness of MPs with template search and model build modes ^46^. The results from the SwissModel server provide a global model quality estimation (GMQE) to evaluate the quality and accuracy of structural models. A score for homology models is provided with interactive 3D viewers.

## Data Availability and Python Libraries

Visualizations and large tables are available on GitHub. https://doi.org/10.5281/zenodo.17170198.

The following Python libraries were utilized in the analysis: (i) Matplotlib – for data visualization and plotting. (ii) SciPy (scipy.stats) – for performing statistical analyses, including the chi-square test. (iii) Statsmodels – for multiple testing correction using the false discovery rate (FDR) approach. (iv) CD-HIT, for UniRef style clustering. Source data for the analysis are provided in the Supplementary materials Supplementary Tables S1-S7.

## Declaration of competing interest

The authors report no conflict of interest.

## Authors contribution

SZ and ML performed the, analysis and visualization. SZ performed the data collection, statistics and produced the code and data for this study. ML, mentoring, conceptualization, writing initial draft. SZ and ML wrote the final draft.

## Funding

This study was supported in part by the Center for Interdisciplinary Data Science Research (CIDR, 3035000440).

## Acknowledgments

We thank the members of Linial’s lab for their support throughout this work, with special thanks to Roei Zucker and Keren Zohar. We thank the GeneCard team (Weizmann Institute of Science) for providing the GeneCaRNA mapping protocol that combined and harmonized many lncRNA resources.

## Supplementary Figures

**Fig. S1.**
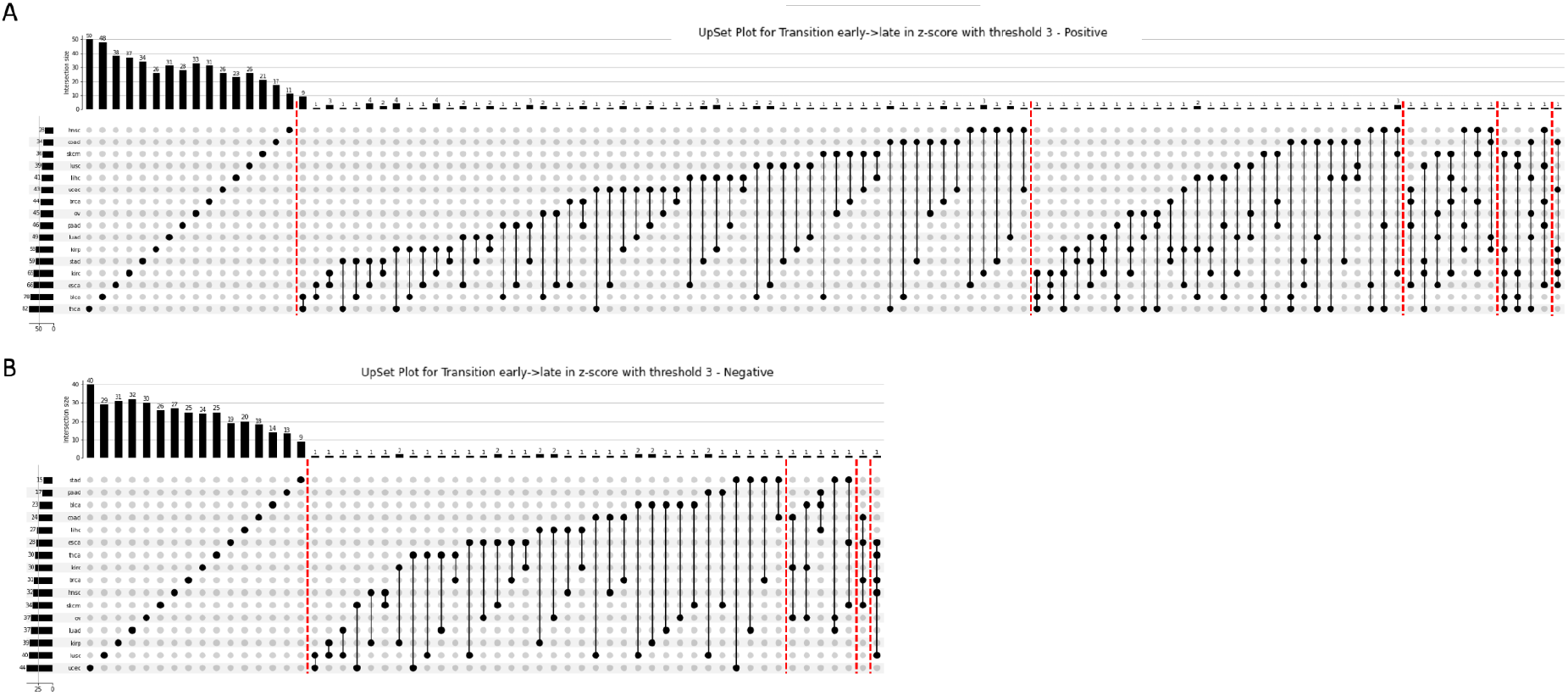
Overlapping Tr-lncRNAs across major cancer types. UpSet plots for the intersection and connectivity of Tr-lncRNAs across major cancer types during early (I & II) to late (III & IV) (E→L) transition. **(A)** Upregulated Tr-lncRNAs (positive) reflect increased expression in the T stage. The dashed red vertical lines group the Tr-lncRNAs by the number of cancer types. **(B)** Downregulated Tr-lncRNAs (negative) reflect decreased expression in the T stage. All combinations of cancer types are shown. The dashed red vertical lines group the Tr-lncRNAs by the number of cancer types. Horizontal bars (left) summarize the total number of significantly changing lncRNAs in each cancer type (for cancer type abbreviations, see Supplemental **Table S1**). Vertical bars (top) count the number of shared lncRNAs across any combination of cancer types.

**Fig. S2.**
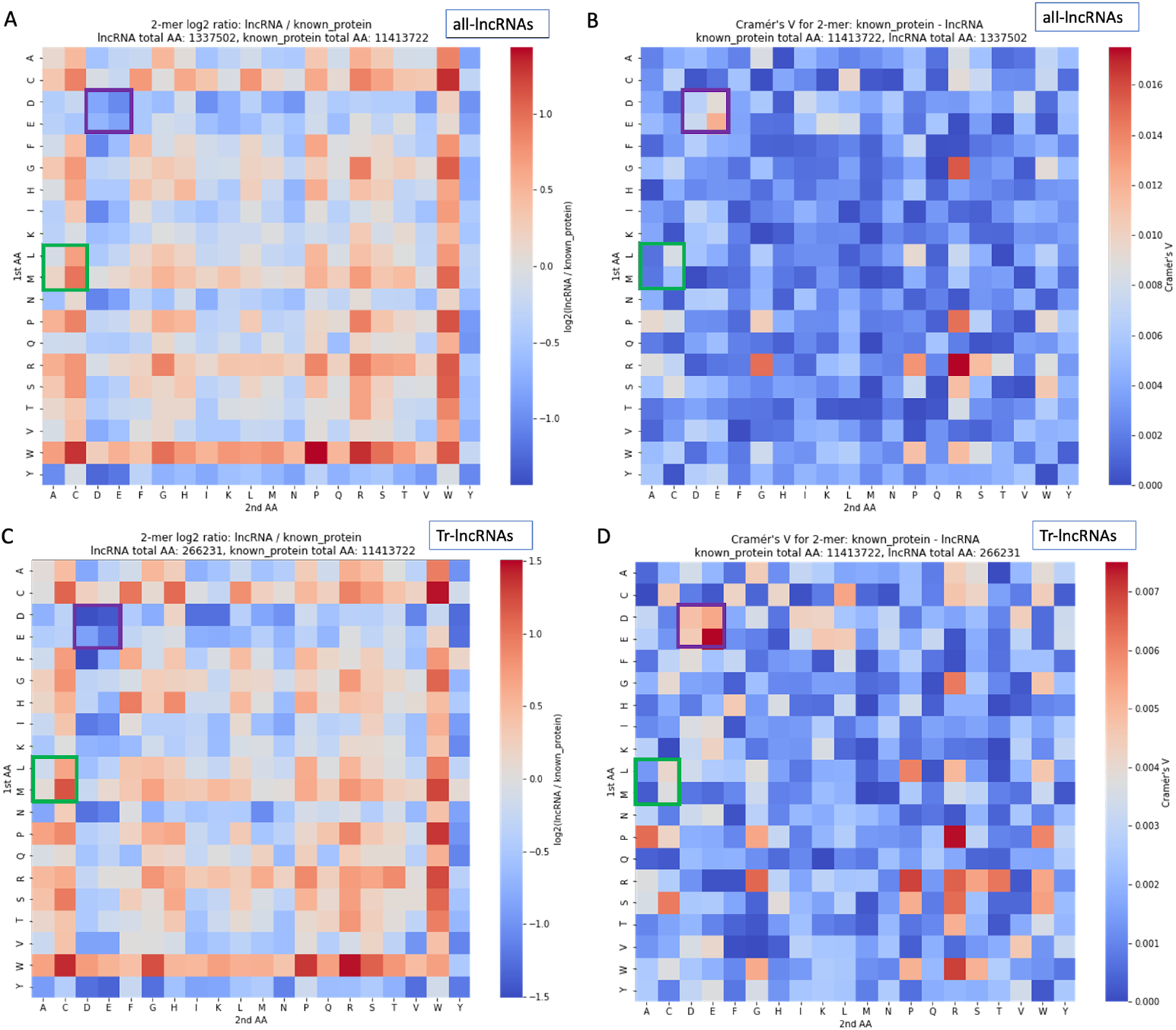
Matrices showing all 400 pairs of dipeptides for unfiltered lncRNA-MPs. **(A)** Heatmap of the log_2_(ratio of lncRNAs to the human proteome). **(B)** A matrix showing the Cramer’s V test for the association of the dipeptides in the human proteome (with 11.44 M aa) vs. the lncRNA-MPs (with 1.34M aa) and the strength of the associations. Values range from 0 to 1.0, where 0 indicates no association and 1.0 indicates maximal association. **(C)** A matrix showing all 400 pairs of dipeptides and their log_2_(ratio of Tr-lncRNAs and human proteome). **(D)** A matrix showing the Cramer’s V test for the association of the dipeptides in the human proteome (with 11.44 M aa) vs. the Tr-lncRNAs (with 266.2k aa) and the strength of the associations. Note that Cramer’s V values are positive, but B and D are scaled individually. The purple and green frames are for visualization purposes to highlight subtle differences in dipeptides across the different analyses.

**Fig. S3.**
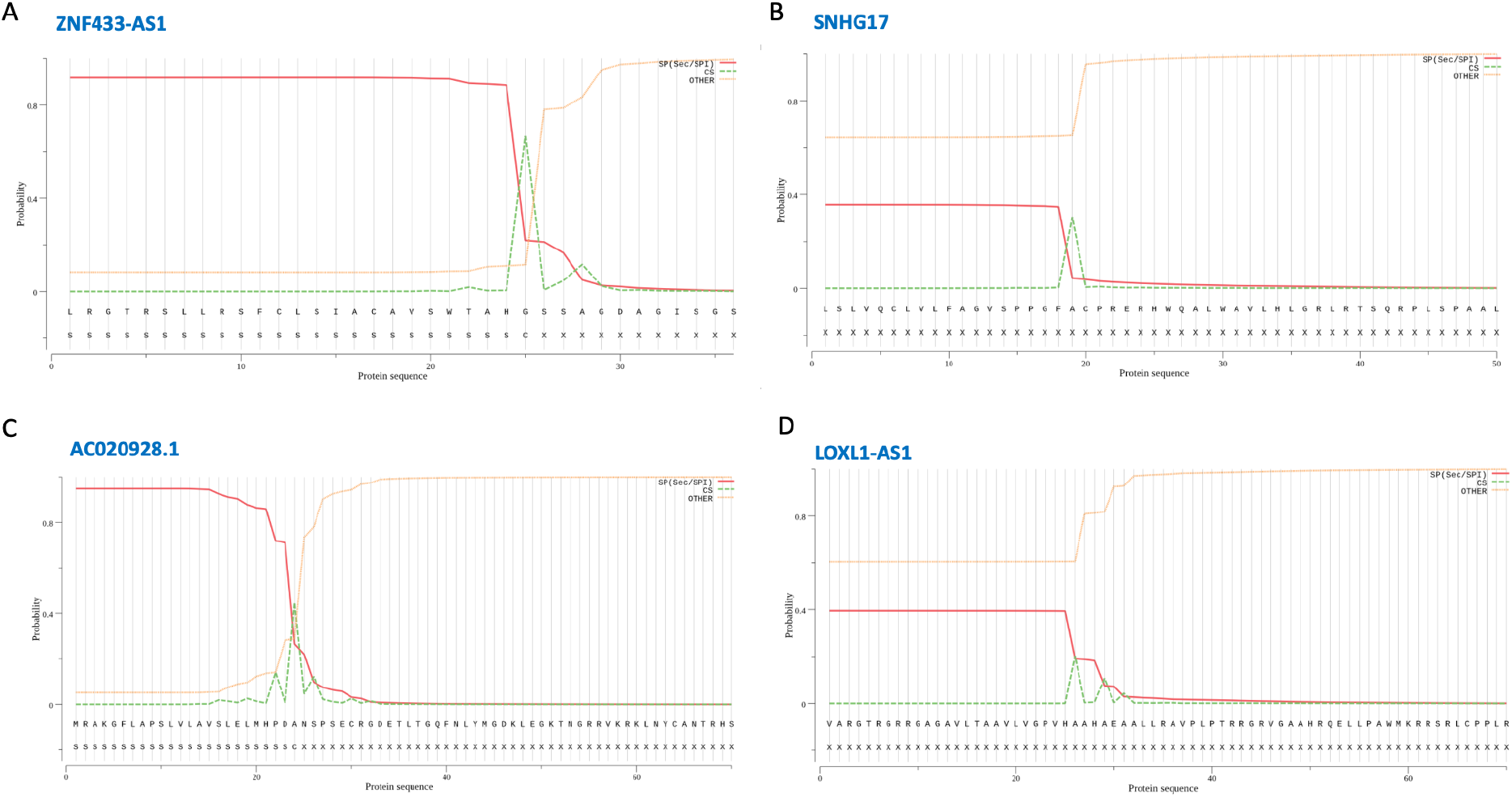
Signal P prediction for secretion potential of lncRNA-encoded MPs. There were 6 sequences (0.034%) that were shown to be a genuine candidate for having a signal peptide (SP). **(A)** ZNF433-AS1. **(B)** AC020928.1. These two genes are predicted with a strong SP prediction, with high Sec/SPI likelihood scores (>0.90). **(C)** SNHG17 and **(D)** LOXL1-AS1 displayed lower SP probabilities (>0.5, SignalP 6.0). CS, cleavage site.

**Fig. S4.**
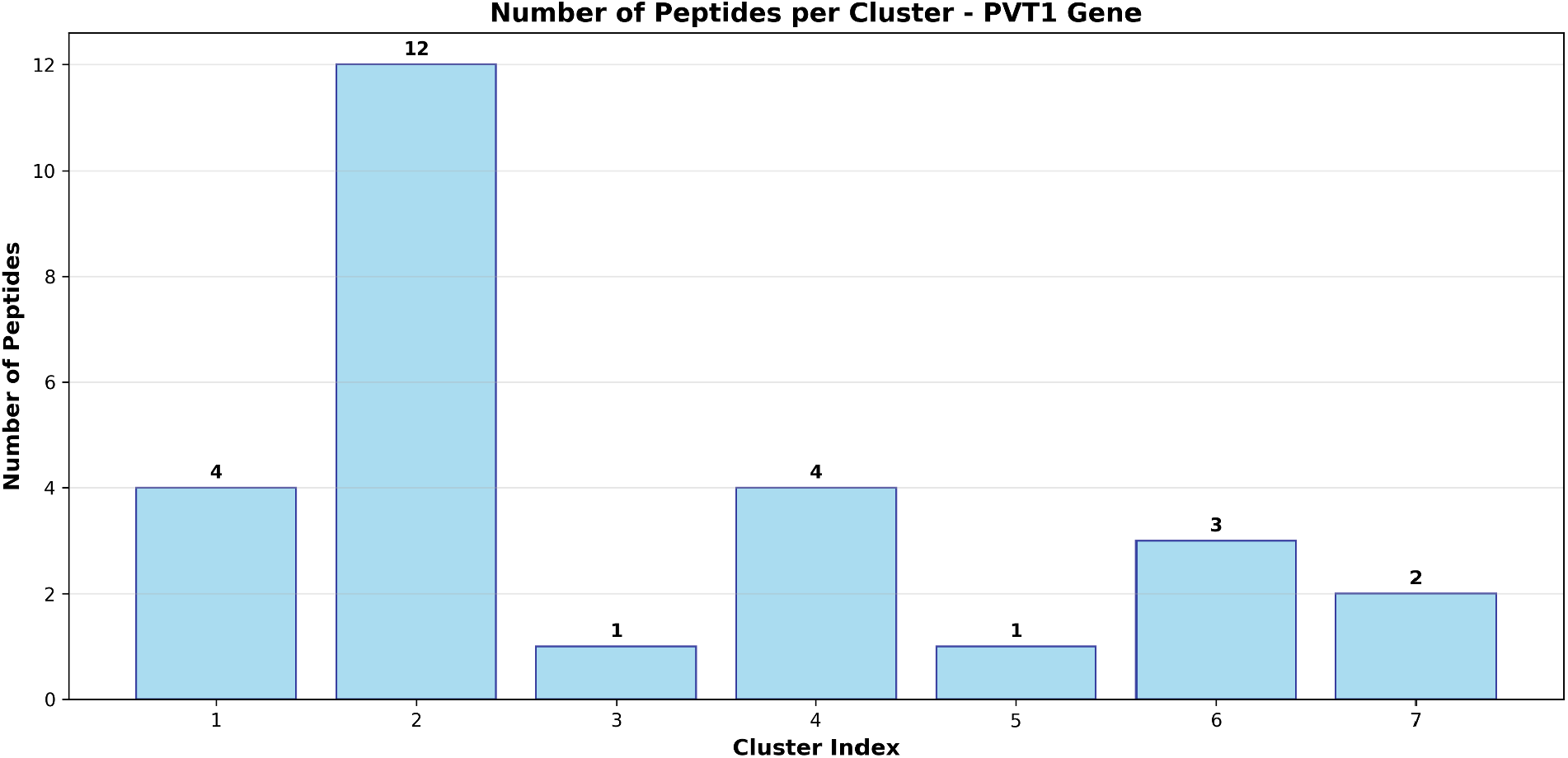
Sequence clustering of PTV1 with 27 high-quality MPs. **(A)** Clustering by sequence identity (>60%) of PTV1 (gene: ENSG00000249859.11). There are 7 peptide sequence-based clusters with non-overlapping sequences across the clusters.

## Notes

https://doi.org/10.5281/zenodo.17170198

### Competing Interest Statement

The authors have declared no competing interest.

